# Foraging behavior, not prey identity, facilitates niche packing in a tropical montane avifauna

**DOI:** 10.64898/2026.06.06.730573

**Authors:** Jacob Drucker, Abhimanyu Lele, Mason Fidino, Dylan Maddox, Sophie Picq, Elisa Bonaccorso, John Bates

## Abstract

Understanding the mechanisms of key ecological and evolutionary patterns and the restructuring of biodiversity in the Anthropocene is contingent on filling knowledge gaps about resource consumption across trophic levels and how resource use is limited by factors intrinsic to organisms and extrinsic aspects of the environment across deep and shallow timespans. We quantified diet composition and foraging behavior across a community of invertivorous birds in the Ecuadorian Andes to explore how resource use facilitates the packing and expansion of niche space across an elevational gradient, contributing the tropical Andes’ status as the most species-rich region on earth. We found evidence that niche packing of morphologically similar species may be offset by greater behavioral plasticity in foraging behavior at species-rich lower elevations where competition is likely more intense and invertebrate prey more diverse. We also tested the extent to which the breadth and similarity of birds’ foraging and dietary niches are shaped by the environmental and competitive gradient across elevation versus species identity and phylogenetic similarity. The specific behaviors and substrates that birds used were far more strongly associated with species identity than elevation, particularly for behaviors requiring specialized morphology that is phylogenetically conserved. In contrast, species identity had little effect on prey selection, which was more strongly associated with elevation. Our findings suggest that elevational range dynamics and niche packing of tropical montane birds are more strongly shaped by phylogenetic constraints on foraging behavior than by specializing on specific prey taxa, highlighting the importance of maintaining structural integrity in tropical forests for preserving functional diversity.

## Introduction

The extent to which the often narrow niches of tropical organisms reflect phylogenetically conserved traits that evolved under climatic stability or are molded by local biotic interactions remains a core question that is key to understanding the evolution and conservation of biodiversity (Dobzhansky 1950; MacArthur 1984; Moles and Ollerton 2016). Stable tropical climates have been associated with the buildup of extreme species richness, resulting in a competitive landscape that is thought to promote the evolution of specialized species and facilitate the packing of niche space (Klopfer and MacArthur 1961; May and MacArthur 1972; Smith et al. 2014; Harvey et al. 2020; Sherry et al. 2020). However, whether specialization to a narrow range of environmental conditions or set of resources limits tropical species ability to respond to anthropogenic-driven global change has become a central question as the climate warms and species and their habitats disappear (Sheldon 2019; Freeman et al. 2021).

Tropical mountains are a nexus for this question because the stacking of species with narrow elevational ranges and the packing of morphologically similar species into niche space along a single elevational gradient makes them global hotspots of species richness and endemism (Cadena et al. 2012; Rahbek et al. 2019). A classic hypothesis explaining narrow elevational distributions in tropical mountains is that limited daily and annual fluctuations in climate have led to reduced climatic niche width relative to temperate latitudes (Huey 1978; Janzen 1967; McCain 2009). Climatic niche specialization has proved a powerful driver in both speciation and community assembly in the Andes and beyond (Cavender-Bares et al. 2011; Bonetti and Wiens 2014; Seeholzer et al. 2017; Bennett et al. 2021), adding credence to a model where species evolve towards an optimal niche in allopatry, disperse linearly within the confines of that climatic envelope, and ultimately come into secondary contact with related species (Endler 1982; Hua 2016). Phylogenetic evidence across the tree of life supports this model, revealing that congeners across a single elevational gradient are not typically each other’s closest relatives (Cavender-Bares et al. 2011; Cadena et al. 2012; Cadena and Céspedes 2020). The implementation of secondary contact in elevational range evolution highlights the role of competition, which can both harden range boundaries between related parapatric species (Jankowski et al. 2013; Freeman 2015) and compress ranges in regions with greater species richness (Freeman et al. 2022).

Regardless of the extent to which the elevational ranges of tropical montane birds and other organisms reflect fundamental or realized niches, it is clear that adaptation to a narrow set of conditions make these species particularly susceptible to shifting geologic features and climates over evolutionary time and to population loss in the face of anthropogenic land use and climate change (García-Moreno et al. 1999; Laurance et al. 2011; Winger and Bates 2015; Freeman et al. 2018; Ausprey et al. 2022). Tropical species are already shifting up mountain slopes faster than their temperate counterparts, and whether they will be able to keep pace with a climate and landscape that is changing rapidly and idiosyncratically remains an urgent question (Linck et al. 2021; Freeman et al. 2021; Colwell and Feeley 2025). Answering this question depends on disentangling the extent to which elevational range shifts reflect species tracking their physiological niche or a set of critical resources, but few studies have addressed resource use explicitly in complex systems such as tropical montane bird communities (Ferger, Schleuning, Hemp, Howell, and Böhning-Gaese 2014; Hanz et al. 2019; Sam et al. 2019; Schumm et al. 2020; Montaño-Centellas et al. 2020; Korejs et al. 2025).

Food is one such resource with a complex relationship to the abiotic environment: temperature, precipitation, primary productivity, and topography can all have profound effects on food availability at all trophic levels and dictate the vegetation structure that enables food acquisition (Davies et al. 2007; Jetz et al. 2009; Ferger, Schleuning, Hemp, Howell, and Böhning-Gaese 2014). Both the quantity and diversity of food resources have been linked to patterns of bird species and functional richness over elevation (Sam et al. 2019; Schumm et al. 2020; Montaño-Centellas et al. 2020; Korejs et al. 2025). Invertebrates are a particularly important resource, reflected in the number of invertivore species that contribute disproportionately to elevated species richness at low and mid-elevations (Pigot et al. 2016; Sam et al. 2019; Schumm et al. 2020; Srinivasan et al. 2024). This swell of bird diversity is not just reflected in the number of species, but also in functional traits that track with the morphological diversity of their prey (Sam et al. 2019; Schumm et al. 2020). That is, a greater variety of sizes and shapes of insects often supports a more functionally diverse avifauna. Furthermore, invertebrates contribute to bird diversity by utilizing different vegetation substrates, like bromeliads and hanging dead leaves, occupying available niche space which consumers can finely partition (Terborgh and Faaborg 1980; Remsen Jr 1985; Marra and Remsen Jr 1997; Sillett et al. 1997). This combination of abundant and diverse invertebrate prey and the vegetation structure they occupy is largely responsible for the observed pattern of niche packing in tropical montane bird communities that facilitates extreme species richness (Pigot et al. 2016; Price et al. 2014; Schumm et al. 2020).

However, quantifying functional traits of avian predators and the biomass and size of their invertebrate prey stops short of explaining how the behaviors of individual birds scale up to broader patterns of resource use that are shaped by the local environment. For example, sympatry of morphologically similar species at lower and mid elevations is common, but how this packing of niche space is facilitated by prey selection and foraging behavior remains unclear (Pigot and Tobias 2015). Foraging behavior offers the ability to reduce competition between morphologically similar species by either specializing on a suite of behaviors to reduce resource overlap, or reducing demand for specific resources by using a greater variety of behaviors (Klopfer and MacArthur 1961; MacArthur and Levins 1967). That is, foraging behavior may vary more where species are more morphologically similar. It is also unclear whether prey selection is a mechanism of niche packing or simply reflects a random sample of the local invertebrate community. If prey selection is random relative to the competitive environment, then dietary niche breadth should be wider at lower elevations and more similar at higher elevations, paralleling the broad decrease in invertebrate diversity (MacArthur 1984; Davies et al. 2007). If prey selection is shaped by competition, then dietary niche breadth should be narrower and more similar at more species-rich lower elevations and broader and less similar at higher elevations.

Despite elevational gradients in climate, competition, and resource availability, how birds access and consume prey resources may ultimately be a function of species-specific traits that are often evolutionarily conserved (Sherry 1990; Montaño-Centellas et al. 2020). More closely-related species are typically more morphologically similar, constraining locomotory and prey handling abilities regardless of their environment (Sherry and McDade 1982). Some predators may also have affinities for specific prey items or foraging substrates that are shared between close relatives (G. H. Rosenberg 1990; K. V. Rosenberg 1990; Sillett et al. 1997). If resource use is phylogenetically conserved, more closely related species should consume the same prey items and use the same foraging behaviors, regardless of the elevation where they live. However, if elevation drives resource use, the probability of using a resource may be similar between species and parallel local abundance of prey items along a mountain slope, particularly if there is strong elevational turnover in prey communities.

We explore the relationship between elevation, species identity, and resource use by quantifying the dietary and foraging niches of invertivorous forest bird species over a 3000 m elevational gradient in the Ecuadorian Andes. We first evaluate the volume and density of niche space across elevation, testing the hypothesis that foraging niche space is more densely packed at lower elevations, parallelling patterns known through morphology. We then quantify the breadth and similarity of foraging and dietary niches between species and communities across elevation, testing for tradeoffs between prey composition and foraging strategies. Finally, we test whether the probability of consuming a prey item or using a foraging behavior is better predicted by elevation, species identity, or phylogenetic similarity. We also explore congruence in turnover between bird communities, arthropod prey communities, and habitat structure across elevation.

## Methods

We quantified dietary and foraging niches using observations of foraging behavior and DNA-metabarcoding of fecal samples collected in the field from invertivorous bird species across a 3000 m elevational gradient in northwestern Ecuador between 2021 and 2024.

### Foraging observations

We collected 2,024 foraging observations by walking trails and roads at field sites within forested habitat (**Figure 1**) and looking for individual birds or mixed species flocks actively searching for food. All observations were collected by JRD (n = 1684) and AL (n= 340) across all hours of the day. Most observations occurred in the context of mixed species flocks. We recorded a single observation per individual bird encountered, which enabled us to consider each foraging event as independent, thereby reducing the chance of pseudoreplication (Ghosh-Harihar and Price 2014). We quantified foraging behavior by recording the occurrence of standardized behaviors (Remsen and Robinson 1990) on a custom-formatted ArcMap Survey123 form (Hennig et al. 2023). These behaviors fell into three categories each representing distinct niche axes: ‘Maneuvers’, ‘Substrates’, and ‘Microhabitat’.

**Figure 1:**
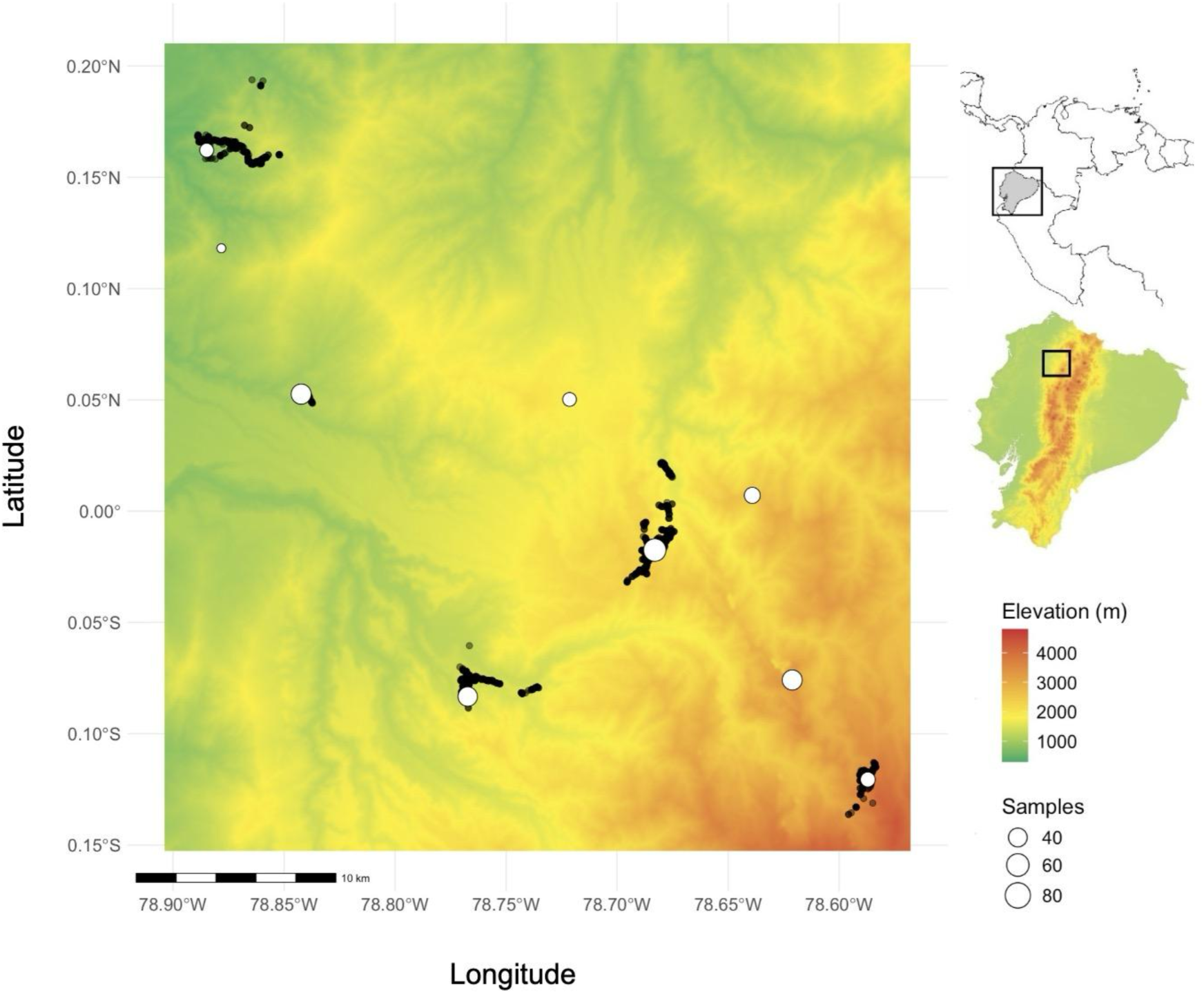
Sampling effort. Map of sampling effort along the west slope of the Andes in Pichincha, Ecuador. White points indicate sites where fecal samples were sourced, sized according to the number of samples from that locality, black points indicate locations of foraging observations.

Maneuvers are the behaviors that a bird uses to access and handle a prey item, for example ‘probing’ or ‘gleaning’. We often recorded more than one maneuver per observation to reflect instances of birds using multiple behaviors to capture prey. For example, ‘hanging’ followed by ‘flaking’, then ‘probing’ to access a prey item would be scored as showing all three of these behaviors as ‘present’. Substrates are the location where the prey item was accessed, such as an epiphytic plant or leaf on a woody plant. Only one substrate was recorded per observation. Microhabitat refers to the environment in an imaginary one-meter-diameter sphere around where the bird attacked a prey item, and is composed of three variables: foliage density, distance from the trunk of the tree, and foraging height. Foliage density and distance from the trunk were scored on a categorical scale of one to five, and height estimations of where the attack occurred relative to the canopy were ultimately binned into five categories from lowest to highest. We compiled our final foraging dataset into a presence-absence matrix of each behavior category for each observation, treating the categorical microhabitat variables as their own columns (e.g. ‘height bin 1’, ‘height bin 2’ etc., are each treated as their own column).

### Fecal Metabarcoding

We collected fecal samples by passively and actively capturing birds with mist nets. We placed extracted birds in clean, wax-lined paper bags until processing within fifteen to thirty minutes. We removed fecal samples from the paper bags using a nylon-based swab or sterilized lab spatula and relocated them to 2 ml tubes filled with 99% ethanol for storage before DNA extraction. Samples were stored at ambient temperature in the field and at -20°C in freezers at the Universidad San Francisco de Quito, Ecuador, and the Field Museum of Natural History, Chicago, USA. Sampling effort for collecting foraging observations and fecal samples was concordant across elevations. We collected fecal samples indiscriminately among captured Passerines but focus on invertivores in this study, processing 480 samples from 22 species (Sherry et al. 2020, Table S1).

Prior to starting batches of DNA extraction, we removed the ethanol and dried the samples by spinning them in a heat-vac before returning them to a -20°C freezer. We extracted DNA from the samples using the Zymo Mini Soil/Fecal kit (Zymo Research Cat. No. D6010), following the manufacturer’s protocol except for using a double elution with 50 μl elution buffer, and vortexing the samples for twenty minutes during the bead-beating step. We extracted samples in batches of eight to twenty-four that were randomized with respect to species and elevation.

We constructed amplicon libraries of the Cytochrome-Oxidase-1 mitochondrial gene with ANML primers (Jusino et al. 2019) to target amplification of invertebrate DNA with iTru compatible tags. We followed the PCR protocols used by (Forsman et al. 2022) with minor modifications. PCR reactions were carried out in 25 µL reactions containing 1× PCR buffer, 2.5 mM MgCl₂, 200 µM each dNTP, 0.2 µM of each primer, and 0.4 U Phusion HF Polymerase. The thermal protocol consisted of an initial denaturation at 94 °C for 5 min, followed by 35 cycles of 94 °C for 30 s, 55 °C for 30 s, and 72 °C for 1 min, with a final extension at 72 °C for 10 min.

The first PCR reaction was cleaned up with 0.65x serapure beads. For PCR2 we used adaptors from Adapterama[DM1] (Glenn et al. 2019), in 25µl reactions containing 1x HiFi Buffer, 0.3 mM of each dNTP, 0.5 µM of each iTru adapter, and 0.5 µl of Kapa HiFi Polymerase. The thermal protocol for PCR2 consisted of 95C for 3m, and then 10 cycles of 95C for 30s, 55C for 30s, 72C for 45s and then a final extension of 72C for 5m. We also cleaned PCR2 with 0.65x beads. We pooled our libraries for sequencing on a single NovaSeq lane at the University of Chicago Sequencing Core.

We processed the resulting 1.4 billion reads of sequencing data in the Qiime2 and R Studio environments (O’Rourke et al. 2020). After importing our sequences and trimming adapters and indexes with the ‘Cutdadapt’ function, we denoised, dereplicated, and removed chimeras with the DADA2 pipeline set to train on the first million reads, which generated ∼18,000 amplicon sequence variants (ASVs). We assigned ASVs to a reference library of CO1 sequences and corresponding taxonomy trimmed to ANML primers designed by (O’Rourke et al. 2020). Our ASVs matched to 243 taxa, yielding a feature Table organized as each sample per row and each column per detected taxon, with the number of reads per prey taxa per sample filling the cells. We considered a prey item ‘present’ if it was represented by more than 300 reads at the Order level or 800 reads at the family level. We used these thresholds based on examining read counts for false/true positive detections of host (bird) taxon recovered in the sample. Our final feature table consisted of presences and absences of only Invertebrate orders and families with manually verified occurrence in the Neotropics.

### Quantifying volume and density of niche space

#### Morphology and foraging behavior

We quantified the volume and density of ordinated morphometric and foraging behavior space across four elevational bins. We identified elevational bins using the FastTopics R package to cluster bird species co-occurring in 50m elevational increments into four community motifs (Dey et al. 2017). Bird occurrences were based on eBird data curated by Freeman et al. (2022) from the Esmeraldas River Basin and verified or modified based on manual review. Elevational breaks for the bird community bins were defined as the elevation where one motif exceeded 50% (**Figure 2**). The resulting elevational bins were 750‒1250m, 1250‒1850m, 1850‒2650m, and 2650‒3850m, corresponding to ‘foothill’, ‘lower cloudforest’, ‘upper cloudforest’, and ‘elfin-treeline’ communities.

**Figure 2:**
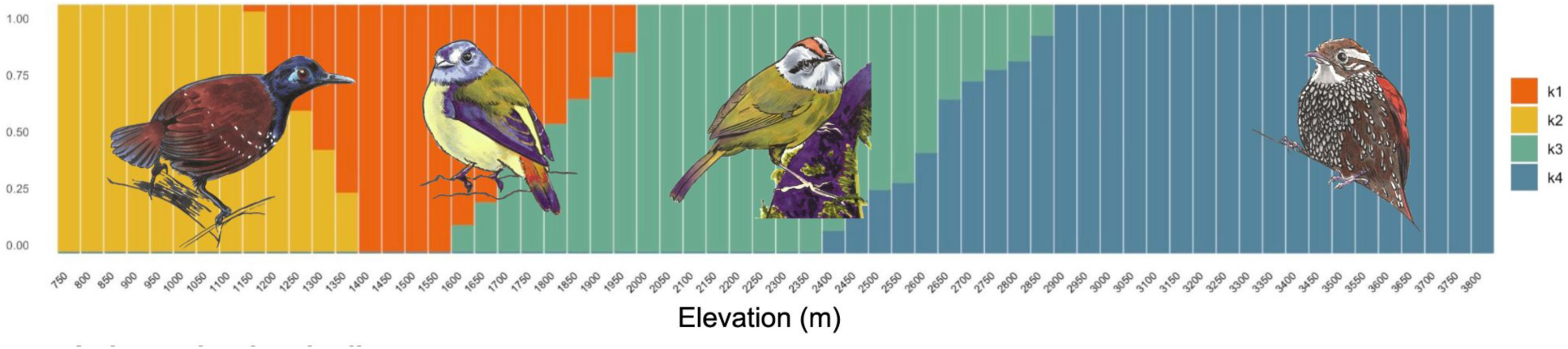
Invertivorous bird community motifs. Turnover in invertivorous bird community motifs based on presence or absence in 50m elevational bins across our full elevational gradient, with motifs identified with the R package FastTopics (Dey et al. 2017). Each bar shows the proportion of each community assigned to each motif. Representative species of each motif are placed accordingly along the gradient. Original art by Alexandra Dube.

We first tested for the occurrence of packing using morphometric methods (Pigot et al. 2016; Pellissier et al. 2018; Crouch et al. 2019; Schumm et al. 2020). We filtered the Avonet morphological dataset (Tobias et al. 2022) to passerine species occurring in our study region (see above, n = 660 species). We scaled morphological traits to a mean of 0 and variance of 1 before using a Principal Component Analysis to quantify species in multivariate space (PC1 = 60% of variance, PC2 = 13% of variance, PC3 = 10% of variance). PC1 corresponded primarily with body size, PC2 with bill length and shape, and PC3 with tail length. We then calculated a convex hull around the species point cloud in 3D PCA space to quantify the volume of morphometric space and the Euclidean nearest neighbor distances between species to quantify the density of morphospace for each elevational bin, interpreting packing as the metric of 1-NND/Hull volume (**Figure 3**).

**Figure 3:**
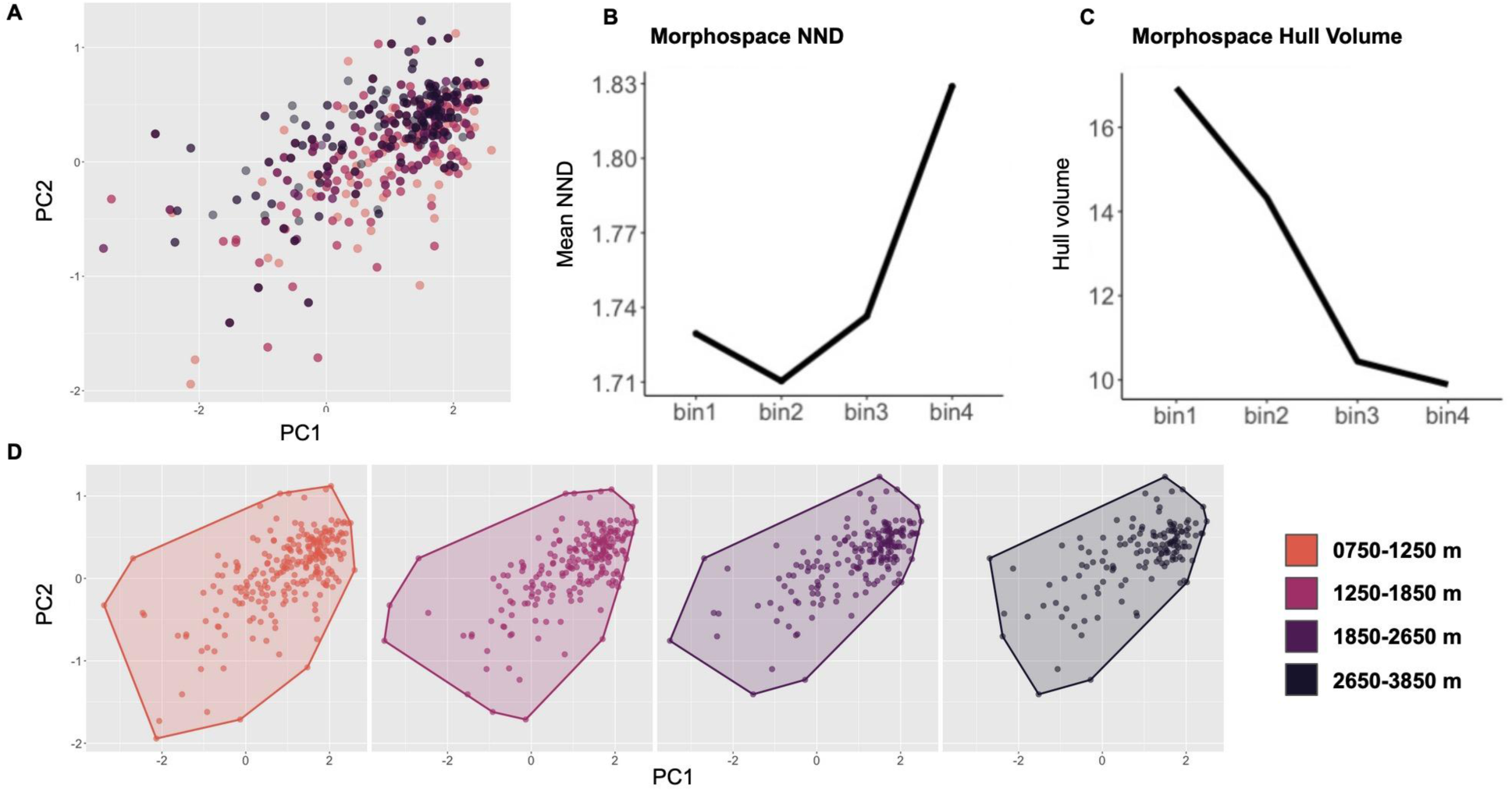
Morphological packing across elevational bins. Components of morphological niche packing along the Pichincha gradient. A) First two axes of a principal component analysis of nine functional traits from all invertivorous passerines regularly occurring in our study region. B) Mean nearest neighbor distance between species in Euclidean space for each elevational bin. C) Convex hull volume values around ordinated species points for bird communities in each elevation bin. D) Visualization of convex hulls in each elevational bin.

We replicated this approach for foraging behavior using a Principal Coordinate Analysis on the observation x behavior presence-absence matrix using the Ape R package (Paradis et al. 2004). The first five PCoA axes captured a total of 54.17% of variance (PCoA1 = 16%, PCoA2 = 13%, PCoA3 = 10%, **Figure 4**, **Figure 5**). The first axis corresponded to attack maneuvers, with Gleaning species loading onto positive values and Probing species onto negative values. The second axis corresponded mostly with vegetation density, with more open microhabitats closer to the trunk loading onto positive values and more closed microhabitats further from the trunk loading onto negative values. We calculated the centroids in PCoA space for each species, again using the metric of 1-NND/Hull volume as a measurement of niche packing in each elevational bin.

**Figure 4:**
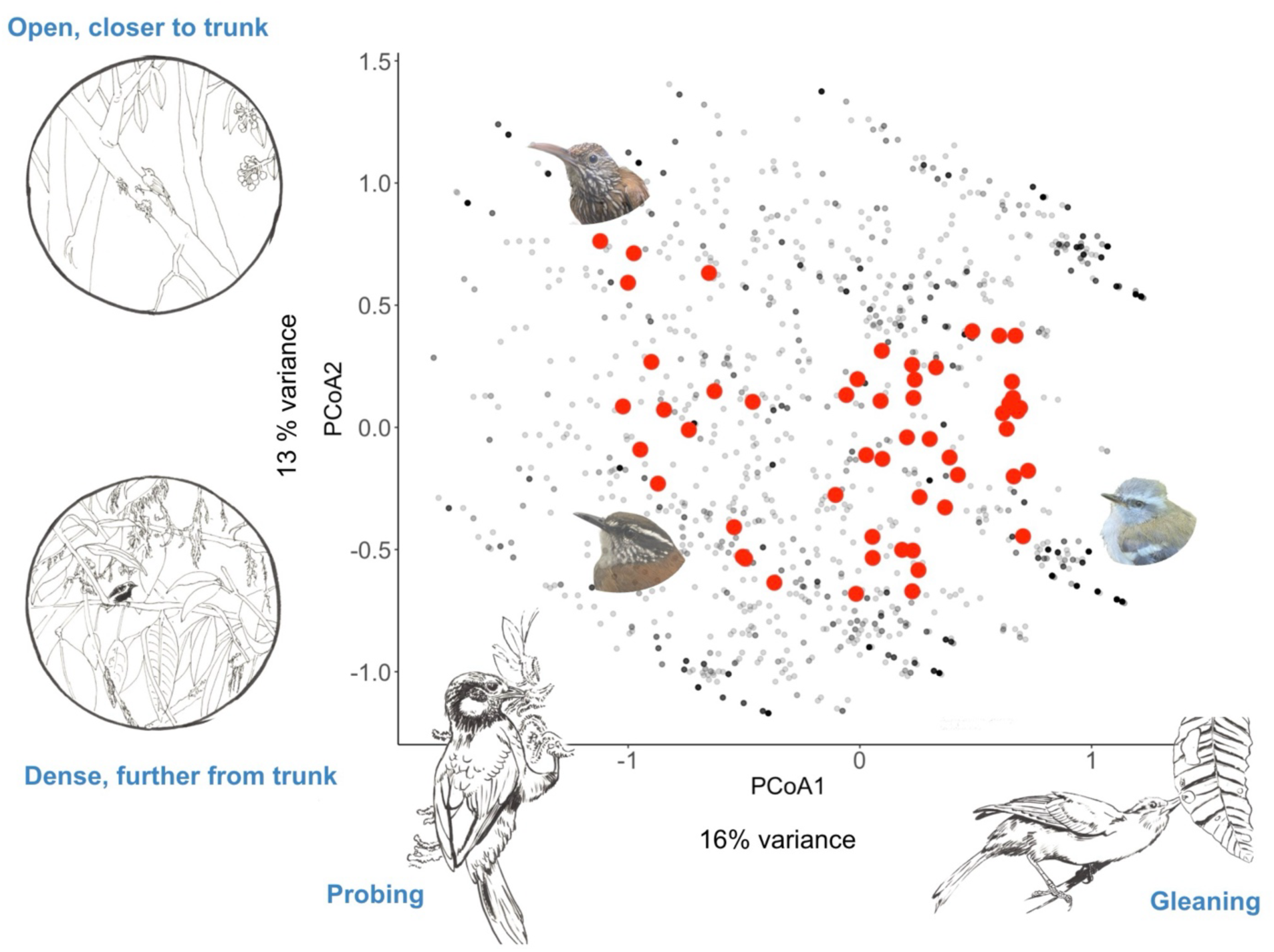
PCoA of foraging niche space. Visualized first two axes of ordinated foraging behavior in a principal coordinate analysis (PCoA) for all observations of all species (grey points), and centroids of species with >= 10 observations (red points). The first axis corresponds largely to maneuver use, with positive values indicating a species’ tendencies to glean more often and negative values indicating a species’ tendency to probe more often. The second axis corresponds mostly to vegetation structure, with higher values indicative of foraging closer to the tree trunk with more open vegetation, and negative values indicative of foraging further from the trunk in denser vegetation. See Figure S1 for visualization of how each behavior loads onto the first twelve axes, which explain 80.85% of total variance.

**Figure 5:**
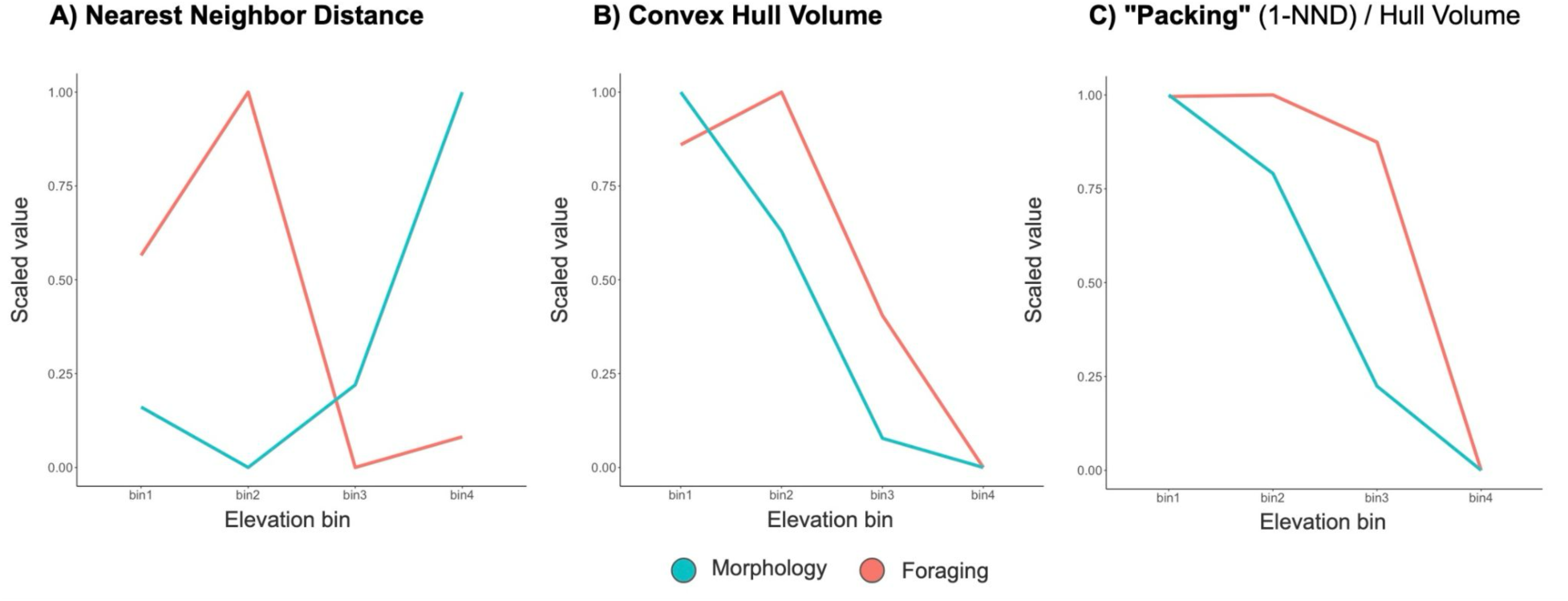
Packing of morphological and foraging niche space. Comparing density and volume of species in ordinated morphometric and foraging behavior space to quantify niche packing across four elevational bins. A) the inverse relationship between species’ morphological and foraging nearest neighbor distances in Euclidean space, B) decrease in volume with elevation but morphospace peaks in foothills while foraging peaks in lower cloud forest, and C) niche packing decreases more evenly in morphospace than foraging space with elevation. Bin 1 = 750 - 1250m, Bin 2 = 1250-1850m, Bin 3 = 1850-2650m, Bin 4 = 2650-3850 m.

#### Prey

We ordinated the prey communities detected in each sample using non-metric multidimensional scaling (NMDS) for invertebrate orders (stress = 0.18) and families (stress = 0.14). Because NMDS is calculated in relative and not Euclidean distances we were unable to use convex hulls and NND values to quantify niche packing. However, we tested for community nestedness between elevational bins using the “nestednodf” function in the Vegan package (Dixon 2003). Higher values of nestedness in one elevational bin indicate that the community in that bin is a subset of the community in another bin. We tested for nestedness separately for invertebrate order and family communities. We also compared counts of the number of invertebrate orders and families detected in each elevational bin, normalizing by the number of observations and mean number of reads in each bin.

### Quantifying niche breadth and composition

We treat dietary niche breadth as the number of prey taxa (orders or families) detected in a sample or species. To quantify foraging niche breadth, we randomly rarefied foraging observations to ten per species with sufficient sample sizes, and calculated the hypervolume around these points, using the global smoothing bandwidth estimated for all two-thousand foraging observations (Blonder et al. 2018).

We compared the similarity of each diet sample or foraging observation to others from the same species using Czekanowski’s proportional similarity index (PSi), implemented with the RInSp R package (Zaccarelli et al. 2013). This metric calculates the frequency of category *i* in individual *j*’s diet relative to the frequency of category *i* in the species as a whole. Higher values of PSi indicate an individual’s resource use is similar to other conspecifics, while negative values indicate an individual uses a different set of resources from other conspecifics.

### Modeling effects of elevation and phylogeny on niche dynamics

We quantified the relationship between elevation, phylogeny, and niche metrics using Bayesian linear mixed effect models in brms (Bürkner 2017). We modeled separately for invertebrate orders and families: 1) the baseline relationship between diet breadth and elevation without species-level effects, using diet breadth as the response variable and the sampling elevation as a single fixed effect, 2) the relationship between diet breadth and elevation within each species, using species’ elevational midpoint as a fixed effect and species identity and phylogenetic covariance as random effects, offset by the logged number of observations for each species, and 3) the relationship between species’ diet and foraging similarity across individuals and midpoint of their elevational range. We modeled species-level foraging niche breadth using only phylogenetic covariance as a random effect given the single value per species, versus aggregated multiple samples in the diet dataset.

We tested whether species identity or elevation better explains diet composition or foraging behavior by modeling the probability of a prey item or behavior’s occurrence in each sample or observation as correlated Bernouli trials in a Bayesian hierarchical mixed effect model (**Table 2**). For these probability models, we used a species’ elevational midpoint as a fixed effect and species and phylogenetic covariance as a random effect, with these random effects considered both independently and jointly. We only modeled the probabilities of prey taxa and foraging behaviors with more than 50 observations.

For all models, elevational fixed effects were scaled to a mean of zero and a variance of one. We planned to include phylogenetic covariance as a random effect in all models. However, in many of our linear models the variance introduced by phylogeny was large enough to obscure the relationship between species dietary niche breadth and elevation, so we proceeded to use only species as a random effect in these models. Furthermore, correlated Bernoulli models with phylogenetic covariance as a random effect struggled to converge in our foraging dataset, so we only modeled behavior correlation when using species as the random effect. We used separate phylogenies for the foraging and diet data sets downloaded using the clootl package (McTavish et al. 2025) from which we calculated covariance between species using the cov2cor function in Ape (Paradis et al. 2004).

Finally, to capture individual level variation in resource use, we estimated the marginal effect of elevation on the probability of using each prey item or foraging behavior by comparing posterior predictions from our fitted models. Specifically, we calculated the difference between the median posterior predicted probability at the mean observed elevation and the median probability at one standard deviation above the mean elevation. We calculated marginal effects separately for each species x behavior combination for the most common prey items and aggregated across species to recover community-wide sensitivity to elevation for each prey category or behavior.

## Results

### Volume and density of niche space across elevational communities

In general, we found evidence of higher-volume, more densely packed niche space at lower elevations based on functional morphology and foraging behavior (**Figure 6**), and expanded dietary niche volume at lower elevations, with nested prey community structure across elevation at the order level (**Figure 7**, **Figure S2**).

**Figure 6:**
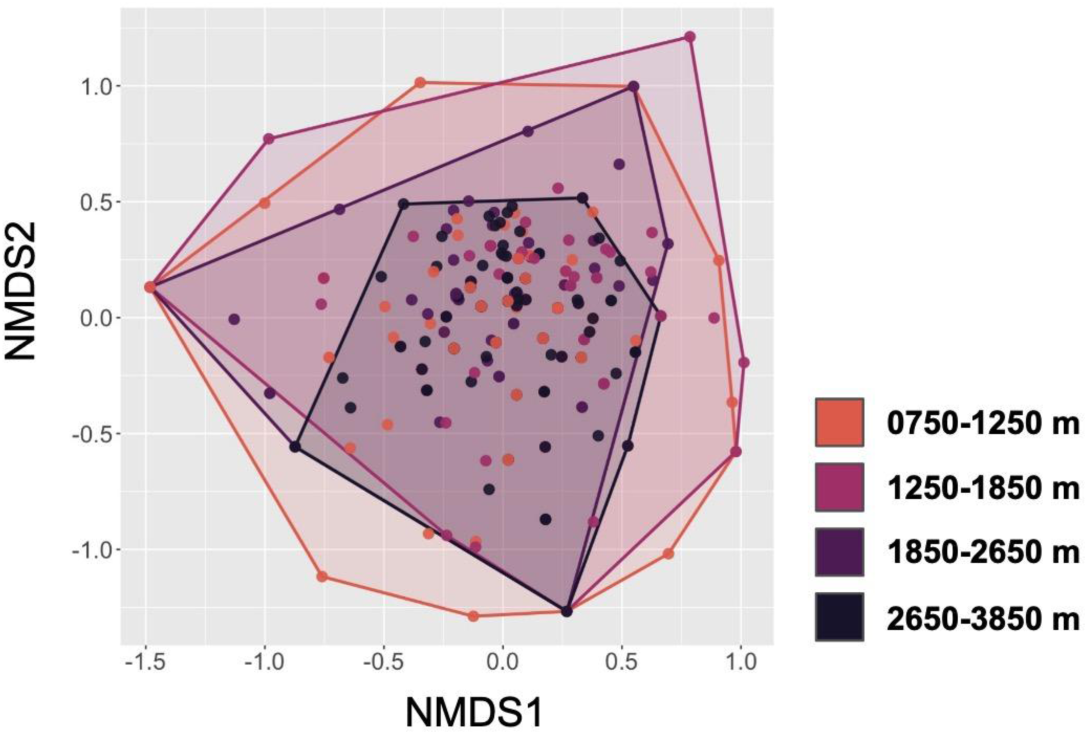
Nested prey orders across elevation bins. Non-metric multidimensional scaling shows that composition of invertebrate prey community orders are nested across elevation, with high-elevation communities representing a subset of low-elevation communities. Each point corresponds to an individual fecal sample, with convex hulls drawn around their extent for each elevational bin to visualize nestedness (NODF = 50.4, Matrix fill = 0.6, Stress = 0.18). See Figures S4 and S5 for species-specific and invertebrate family-level NMD plots.

**Figure 7:**
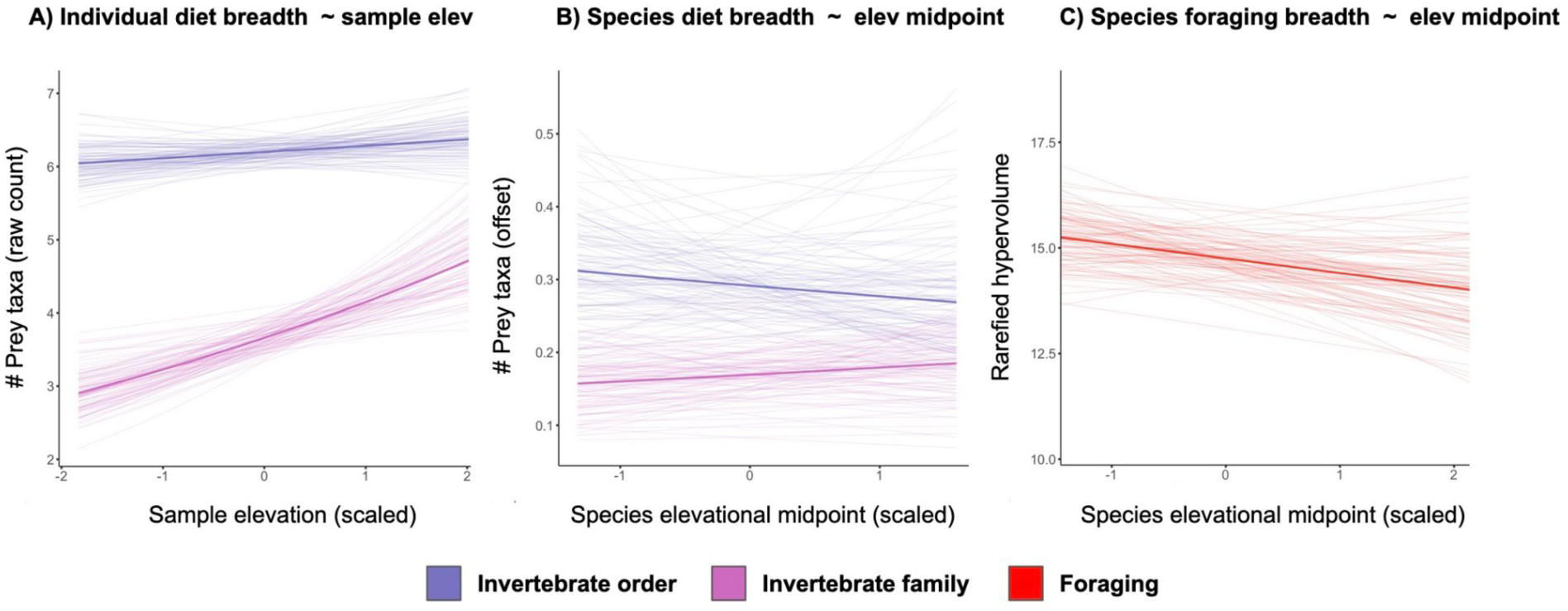
Diet and foraging niche breadth across elevation. Panels show the relationship between elevation and niche breadth based on Bayesian linear mixed effect models for A) diet breadth of an individual without considering species-level effects, B) diet breadth of a species as a function of the midpoint of its elevational range while offsetting for variation in sample size, and C) foraging behavior niche breadth as a function of a species’ elevational midpoint. Bold smooths show the mean posterior estimate of the slope and intercept for each model, with narrow smooths showing variation from 100 random posterior draws. Blue lines represent diet breadth of invertebrate orders, pink lines diet breadth of invertebrate families, and red lines foraging niche breadth.

Hull volumes around ordinated species means in morphological and foraging space decreased sharply at higher elevations. Morphological hull volumes were largest in lower cloud forests, while foraging hull volumes were largest in the foothills. Euclidean nearest-neighbor distances between species in ordinated space showed an inverse relationship between morphology and foraging behavior (**Figure 6**): species were more morphologically similar to each other in lower-elevation communities, and more disparate at higher elevations. However, based on nearest neighbor distances between centroids, species foraged more differently from each other at low elevations and more similarly at high elevations. This contrasting trend of nearest neighbor distances coupled with similarly decreasing Hull volumes across elevations underlies a pattern of more tightly packed foraging niche space along a greater proportion of the elevational gradient than morphological niche space, which gradually becomes less dense at higher elevations (**Figure 7**).

Hull volumes calculated around invertebrate prey communities derived from fecal metabarcoding also decreased at higher elevations at the order level (Figure S2). Rather than occupying different regions of NMDS space in each elevational bin, prey communities were nested across the gradient (*N* species = 50.2, *N* elev bins = 70.9, NODF = 50.4, Matrix fill = 0.63). That is, invertebrate order communities at high elevations were a subset of order communities at lower elevations. At the family level, invertebrate communities occupied similar regions and volumes of NMDS space, showing a low degree of nestedness (Figure S3), *N* species = 36.4, *N* elev. bins = 59.7, NODF = 36.46, Matrix fill = 0.45).

Raw counts of invertebrate orders and families were remarkably similar across elevational bins with order counts ranging between 18 and 21 and family counts between 53 and 64. Taxonomic richness across elevation remained similar when correcting for sampling effort. However, correcting taxa counts by the mean number of reads in each bin showed the highest richness at the lowest elevations, decreasing gradually for orders but remaining similar for families (Figure S4).

### Niche breadth and similarity across elevation

We recovered a generally weak positive relationship between diet breadth and elevation, with moderate differences between species regardless of phylogenetic similarity (**Figure 5**, **Table 1**). Across the predator community (i.e. without considering the effects of species or phylogenetic covariation), diet breadth had a weakly positive relationship with elevation, particularly at the family level (Estimate = 0.13, 95% CIs = 0.00.21, **Figure 5**). Bird species living at higher elevations also showed the trend of consuming more invertebrate families than lower elevation bird species, but with weaker statistical support (Estimate = 0.04, 95% CIs = - 0.2-0.28). The relationship between diet breadth and elevation was more ambiguous at the invertebrate order level, with the mean posterior estimate suggesting lower-elevation predator species have broader dietary niches, although with wide uncertainty (Estimate = -0.03, 95% Cis = -0.26, 0.11).

**Table 1:**
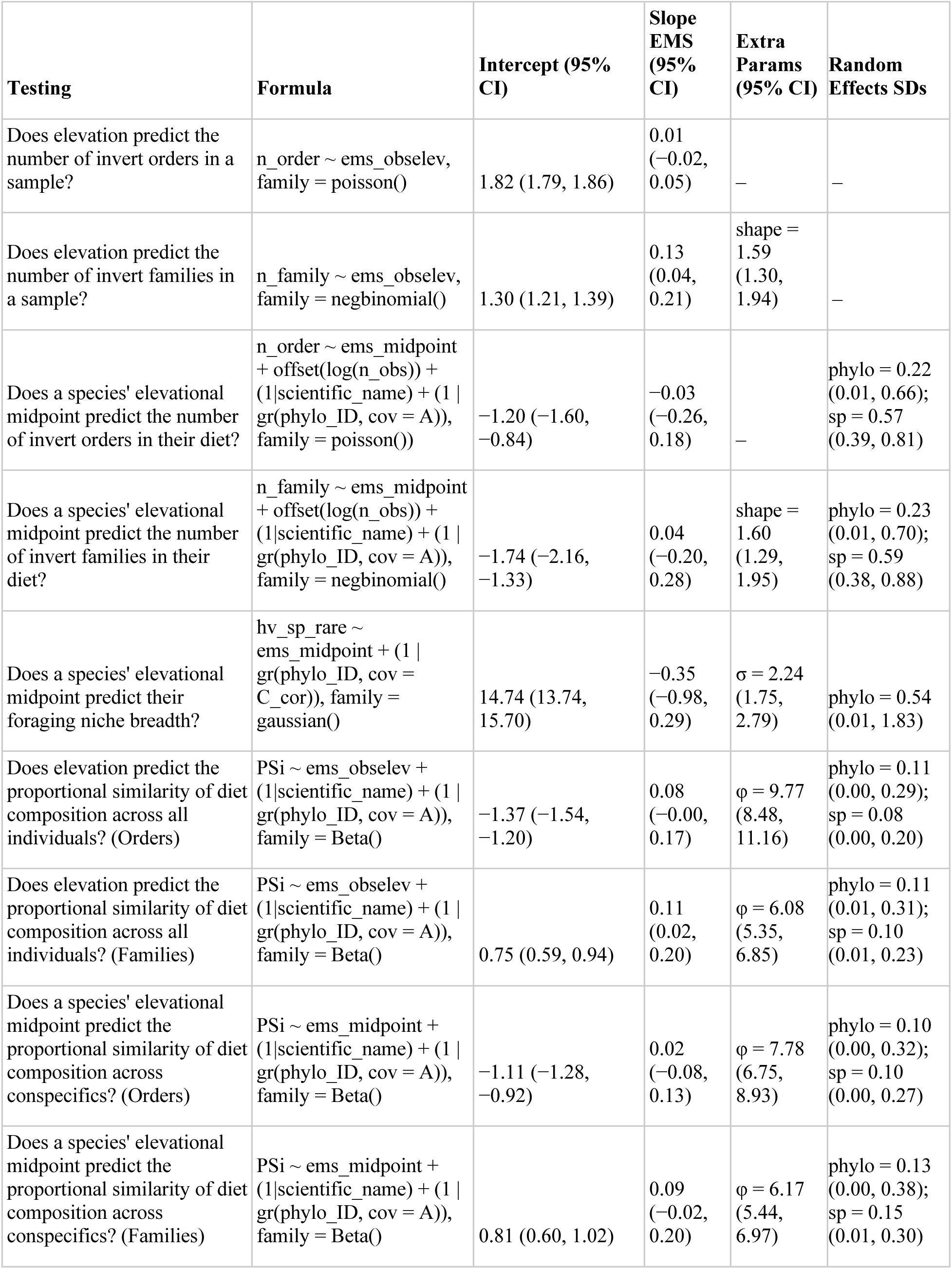
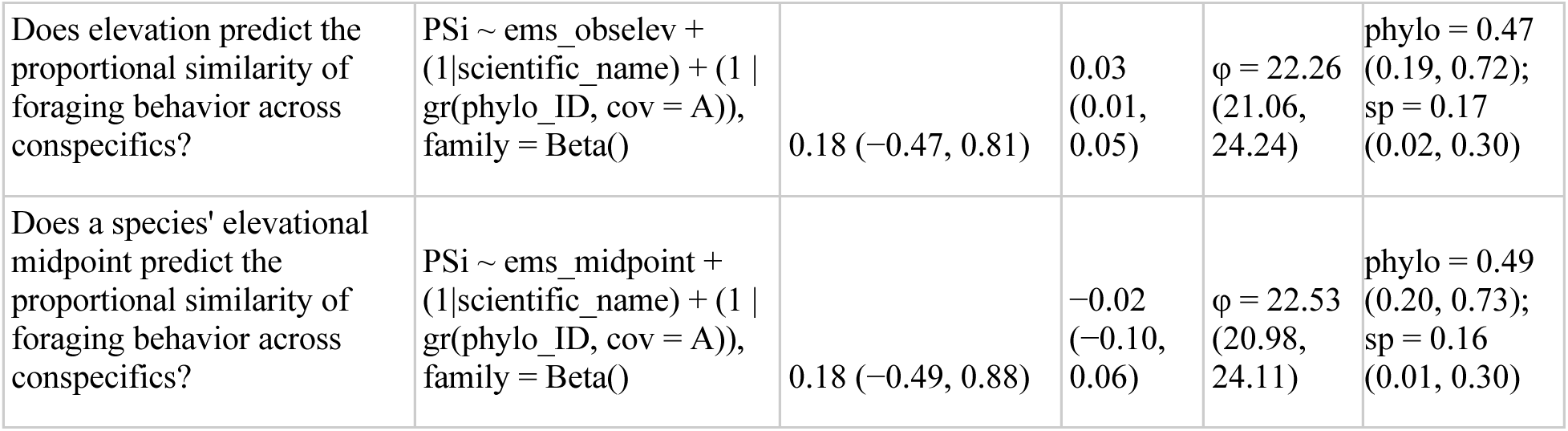
Bayesian linear mixed efffects models of foraging breadth and similarity.

| Testing | Formula | Intercept (95% CI) | Slope EMS (95% CI) | Extra Params (95% CI) | Random Effects SDs |
| --- | --- | --- | --- | --- | --- |
| Does elevation predict the number of invert orders in a sample? | n_order ~ ems_obselev, family = poisson() | 1.82 (1.79, 1.86) | 0.01 (-0.02, 0.05) | – | – |
| Does elevation predict the number of invert families in a sample? | n_family ~ ems_obselev, family = negbinomial() | 1.30 (1.21, 1.39) | 0.13 (0.04, 0.21) | shape = 1.59 (1.30, 1.94) | – |
| Does a species' elevational midpoint predict the number of invert orders in their diet? | n_order ~ ems_midpoint + offset(log(n_obs)) + (1\|scientific_name) + (1\|gr(phylo_ID, cov = A)), family = poisson() | -1.20 (-1.60, -0.84) | -0.03 (-0.26, 0.18) | – | phylo = 0.22 (0.01, 0.66); sp = 0.57 (0.39, 0.81) |
| Does a species' elevational midpoint predict the number of invert families in their diet? | n_family ~ ems_midpoint + offset(log(n_obs)) + (1\|scientific_name) + (1\|gr(phylo_ID, cov = A)), family = negbinomial() | -1.74 (-2.16, -1.33) | 0.04 (-0.20, 0.28) | shape = 1.60 (1.29, 1.95) | phylo = 0.23 (0.01, 0.70); sp = 0.59 (0.38, 0.88) |
| Does a species' elevational midpoint predict their foraging niche breadth? | hv_sp_rare ~ ems_midpoint + (1\|gr(phylo_ID, cov = C_cor)), family = gaussian() | 14.74 (13.74, 15.70) | -0.35 (-0.98, 0.29) | $\sigma$ = 2.24 (1.75, 2.79) | phylo = 0.54 (0.01, 1.83) |
| Does elevation predict the proportional similarity of diet composition across all individuals? (Orders) | PSi ~ ems_obselev + (1\|scientific_name) + (1\|gr(phylo_ID, cov = A)), family = Beta() | -1.37 (-1.54, -1.20) | 0.08 (-0.00, 0.17) | $\phi$ = 9.77 (8.48, 11.16) | phylo = 0.11 (0.00, 0.29); sp = 0.08 (0.00, 0.20) |
| Does elevation predict the proportional similarity of diet composition across all individuals? (Families) | PSi ~ ems_obselev + (1\|scientific_name) + (1\|gr(phylo_ID, cov = A)), family = Beta() | 0.75 (0.59, 0.94) | 0.11 (0.02, 0.20) | $\phi$ = 6.08 (5.35, 6.85) | phylo = 0.11 (0.01, 0.31); sp = 0.10 (0.01, 0.23) |
| Does a species' elevational midpoint predict the proportional similarity of diet composition across conspecifics? (Orders) | PSi ~ ems_midpoint + (1\|scientific_name) + (1\|gr(phylo_ID, cov = A)), family = Beta() | -1.11 (-1.28, -0.92) | 0.02 (-0.08, 0.13) | $\phi$ = 7.78 (6.75, 8.93) | phylo = 0.10 (0.00, 0.32); sp = 0.10 (0.00, 0.27) |
| Does a species' elevational midpoint predict the proportional similarity of diet composition across conspecifics? (Families) | PSi ~ ems_midpoint + (1\|scientific_name) + (1\|gr(phylo_ID, cov = A)), family = Beta() | 0.81 (0.60, 1.02) | 0.09 (-0.02, 0.20) | $\phi$ = 6.17 (5.44, 6.97) | phylo = 0.13 (0.00, 0.38); sp = 0.15 (0.01, 0.30) |
| Does elevation predict the proportional similarity of foraging behavior across conspecifics? | PSi ~ ems_obselev + (1\|scientific_name) + (1 \| gr(phylo_ID, cov = A)), family = Beta() | 0.18 (−0.47, 0.81) | 0.03 (0.01, 0.05) | φ = 22.26 (21.06, 24.24) | phylo = 0.47 (0.19, 0.72);<br>sp = 0.17 (0.02, 0.30) |
| Does a species' elevational midpoint predict the proportional similarity of foraging behavior across conspecifics? | PSi ~ ems_midpoint + (1\|scientific_name) + (1 \| gr(phylo_ID, cov = A)), family = Beta() | 0.18 (−0.49, 0.88) | −0.02 (−0.10, 0.06) | φ = 22.53 (20.98, 24.11) | phylo = 0.49 (0.20, 0.73);<br>sp = 0.16 (0.01, 0.30) |

Rarefied hypervolumes representing species’ foraging niche breadth also showed a negative but uncertain relationship with elevation (**Figure 5c**, **Table 1**, Estimate = -0.35, 95% Cis = -0.98, 0.29). Phylogeny explained some of additional variance in foraging niche breadth (SD = 0.54, 95% CIs = 0.01-1.83) but more variance was explained by residual error (σ = 2.24, 95% CIs= 1.75–2.79).

Diet composition of invertebrate orders and families became more similar at higher elevations at the level of individuals and species, while foraging behavior similarity showed a weak relationship with elevation (**Figure 6**, **Table 1**). Diet similarity showed a clearer relationship with elevation than any other variable, with a simple plot of average intraspecific similarity capturing the dominant pattern in our models (**Figure 7**). Species and phylogenetic covariance both explained relatively little variance in diet similarity but added a significant degree of uncertainty to estimates of foraging similarity (**Table 1**).

### Resource use across elevation

#### Community-wide probabilities of use

We detected 29 invertebrate orders and 105 invertebrate families across 466 samples from 22 bird species (**Figure 8**, Figure S2). Across all species, seven orders occurred in at least 50% of samples, with Diptera and Lepidoptera being the most common. Geometridae (Lepidoptera) and Anyphaenidae (Araneae) were the only families detected in more than 25% of samples (**Table 2**).

**Figure 8:**
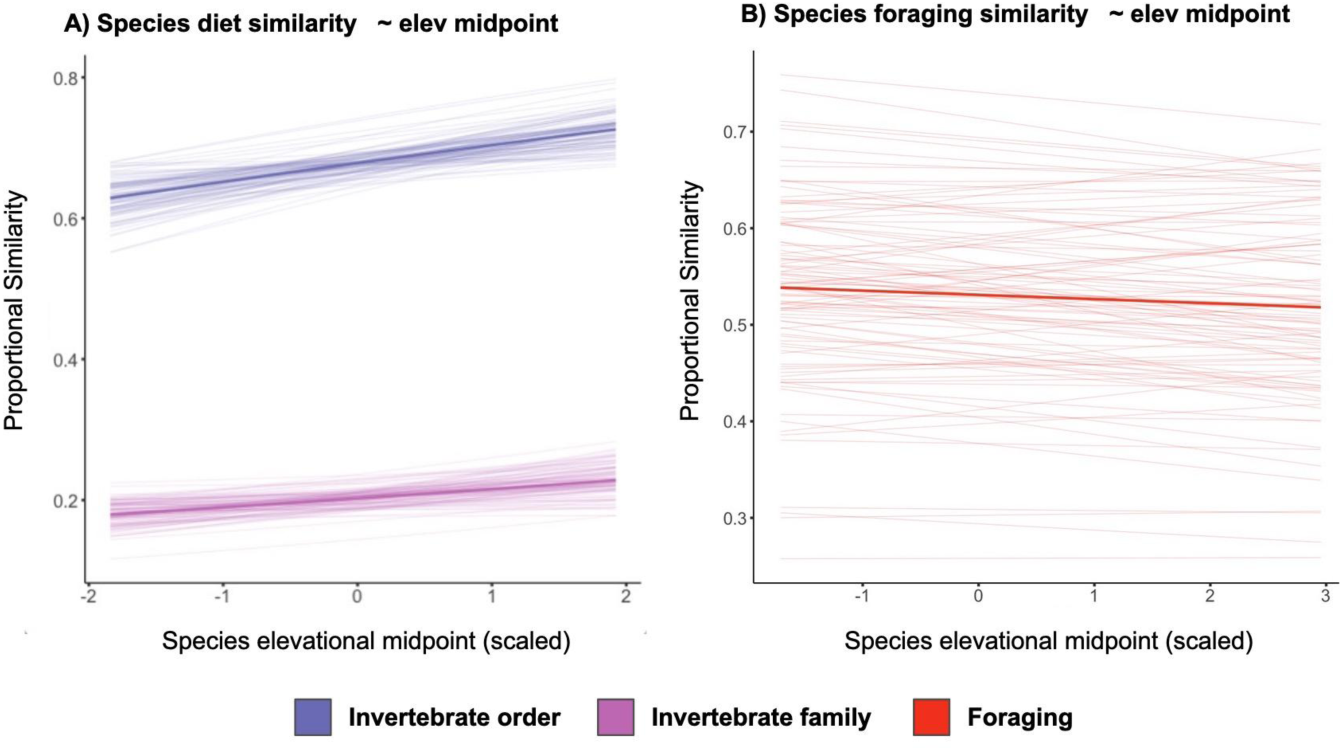
Diet and foraging niche similarity across elevation. Panels show the relationship between elevation and niche similarity between conspecific individuals based on Bayesian linear mixed effect models for A) diet and B) foraging behavior, accounting for phylogenetic covariance and differences in sample sizes between species. Bold smooths show the mean posterior estimate of the slope and intercept for each model, with narrow smooths showing variation from 100 random posterior draws. Blue lines represent diet similarity of invertebrate orders, pink lines diet similarity of invertebrate families, and red lines foraging niche similarity.

**Table 2:**
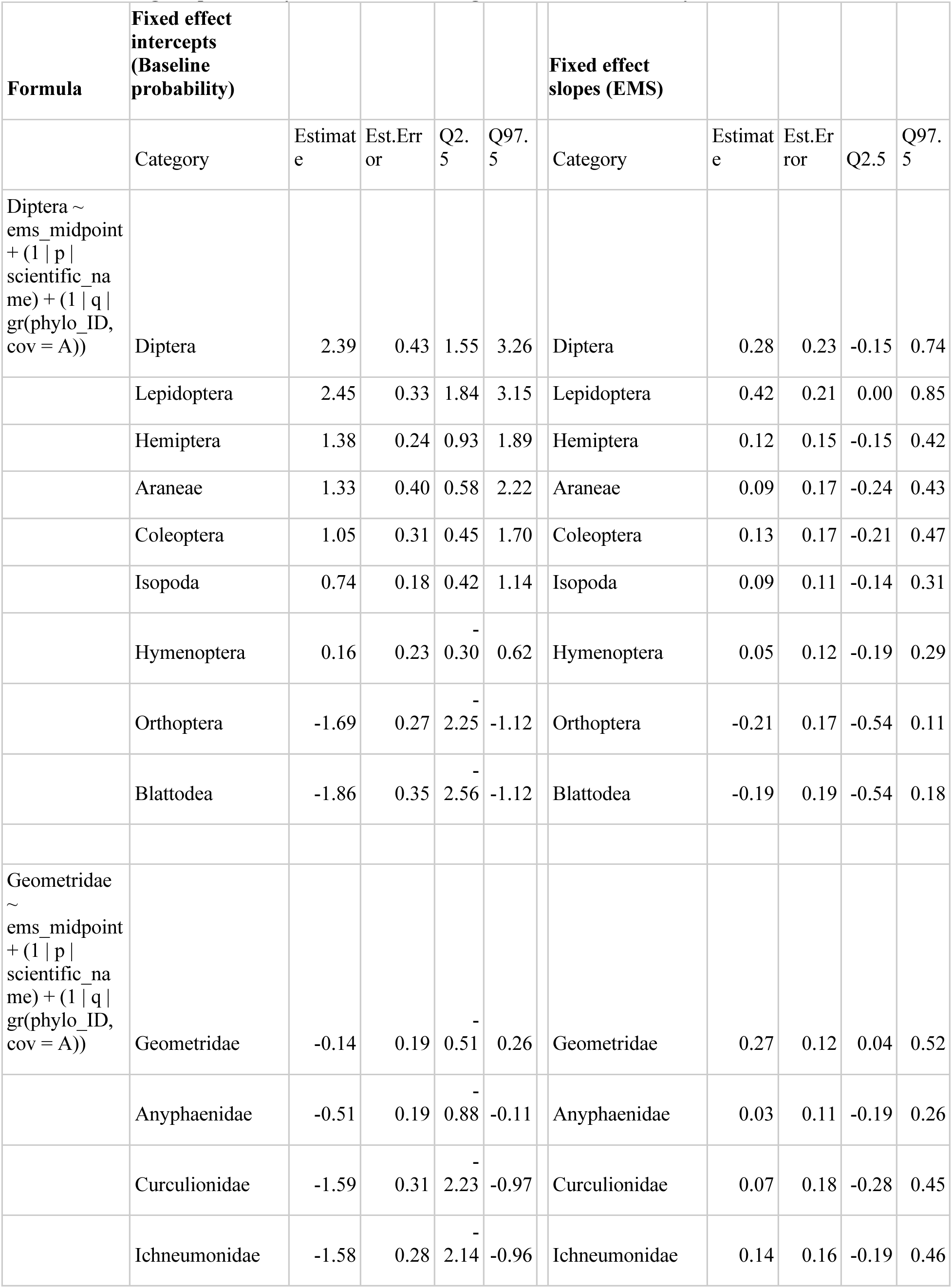
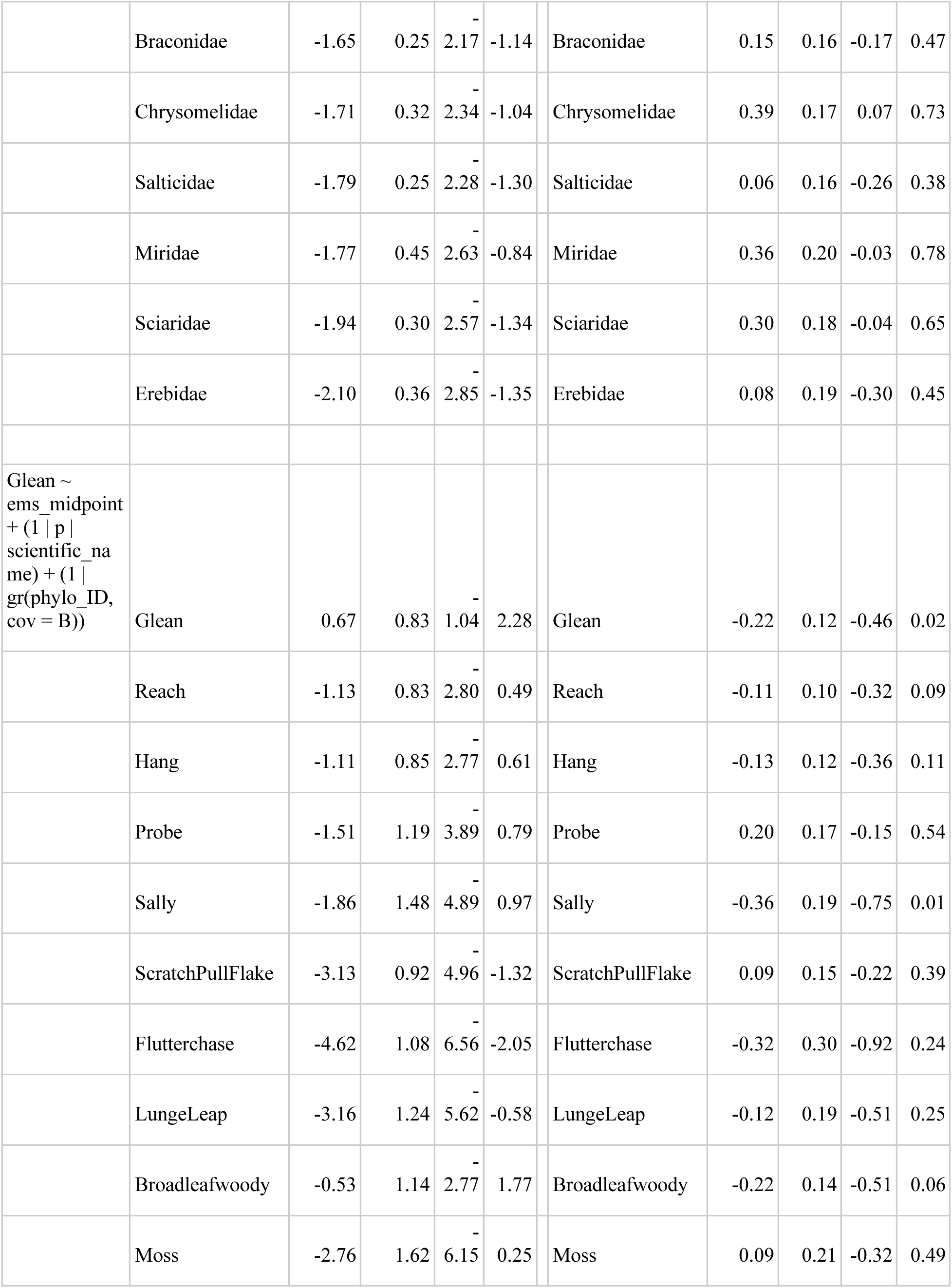
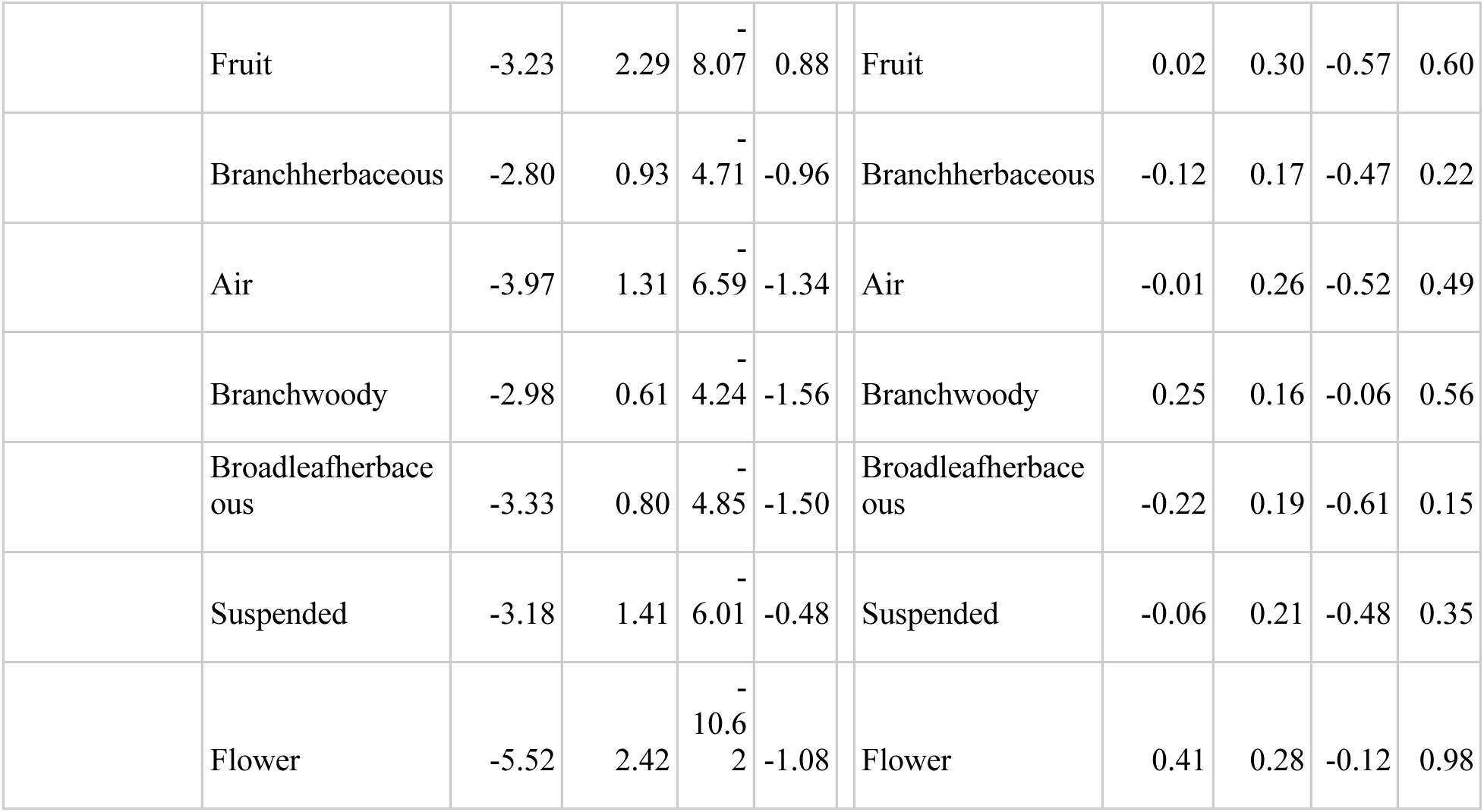
Modeling the probability of resource use using Bernoulli trials in a Bayesian mixed effect model.

Across 1934 foraging observations of 154 bird species, “gleaning” was by far the most common behavior (75% of observations) and “broadleaf woody” the most common substrate (23% of observations). “Flutter chase” was the rarest behavior (< 1% of observations) and “Flower” was the least frequently used substrate (< 1% of observations). Species-level estimates of prey consumption probabilities rarely deviated beyond community-wide means, in contrast with species-level estimates of foraging behavior probability which deviated widely from community-level means (**Figure 9**).

**Figure 9:**
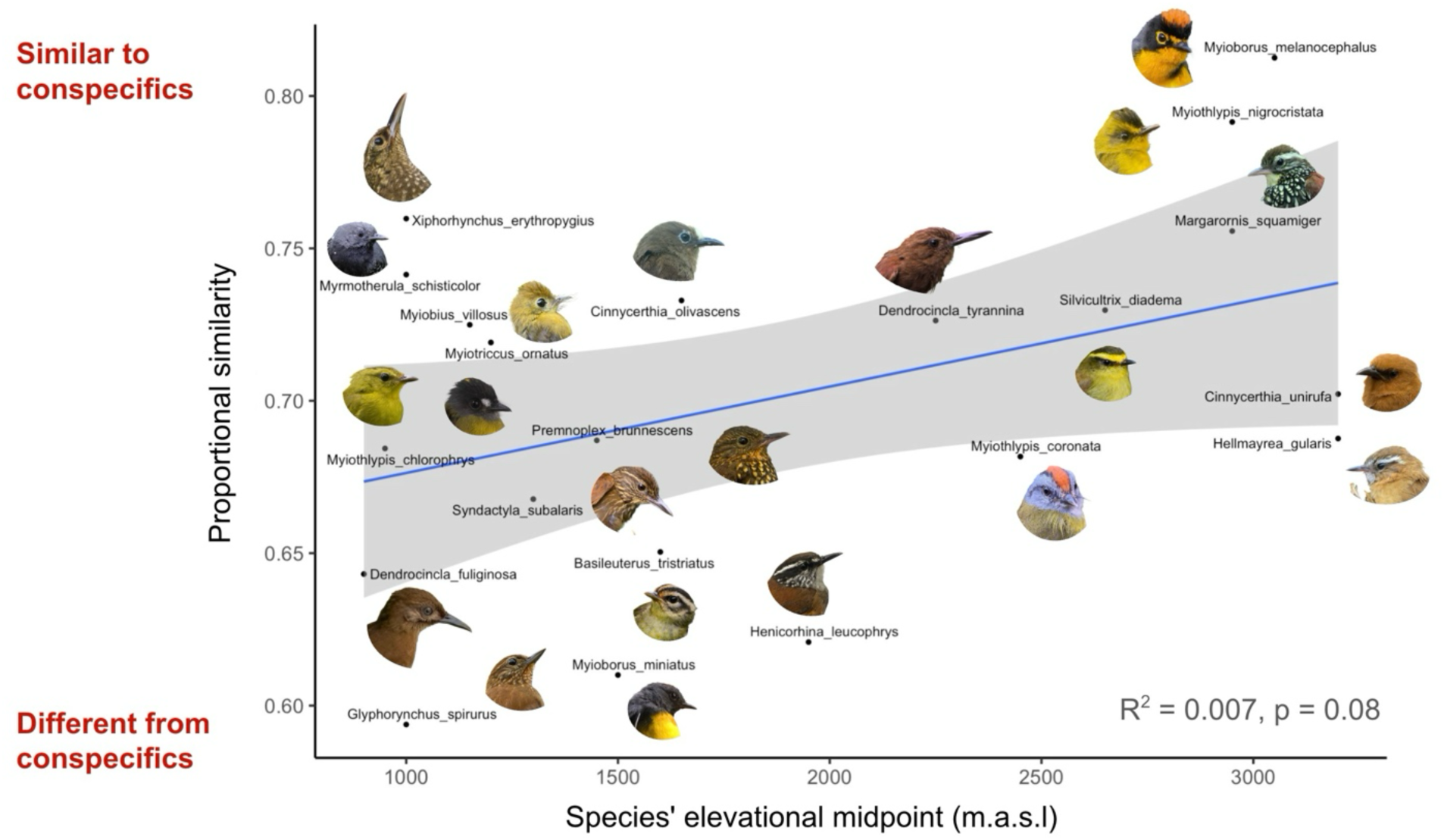
Conspecific diet composition is more simlar among higher-elevation species. Average diet similarity (Czekanowski’s proportional similarity index, Zaccareli et al. 2013) across conspecific fecal samples plotted against the midpoint of the predator species’ elevational range. Blue line is an overlaid slope and intercept from a linear model with 95% confidence intervals in gray.

#### Effect of elevation on probability of behavior or resource use

Elevation generally had a weak effect on the probability of a species performing a foraging behavior or using a resource, but several maneuvers, substrates, or prey taxa were more likely to occur at higher or lower elevations (**Figure 10**, **Table 2**).

**Figure 10:**
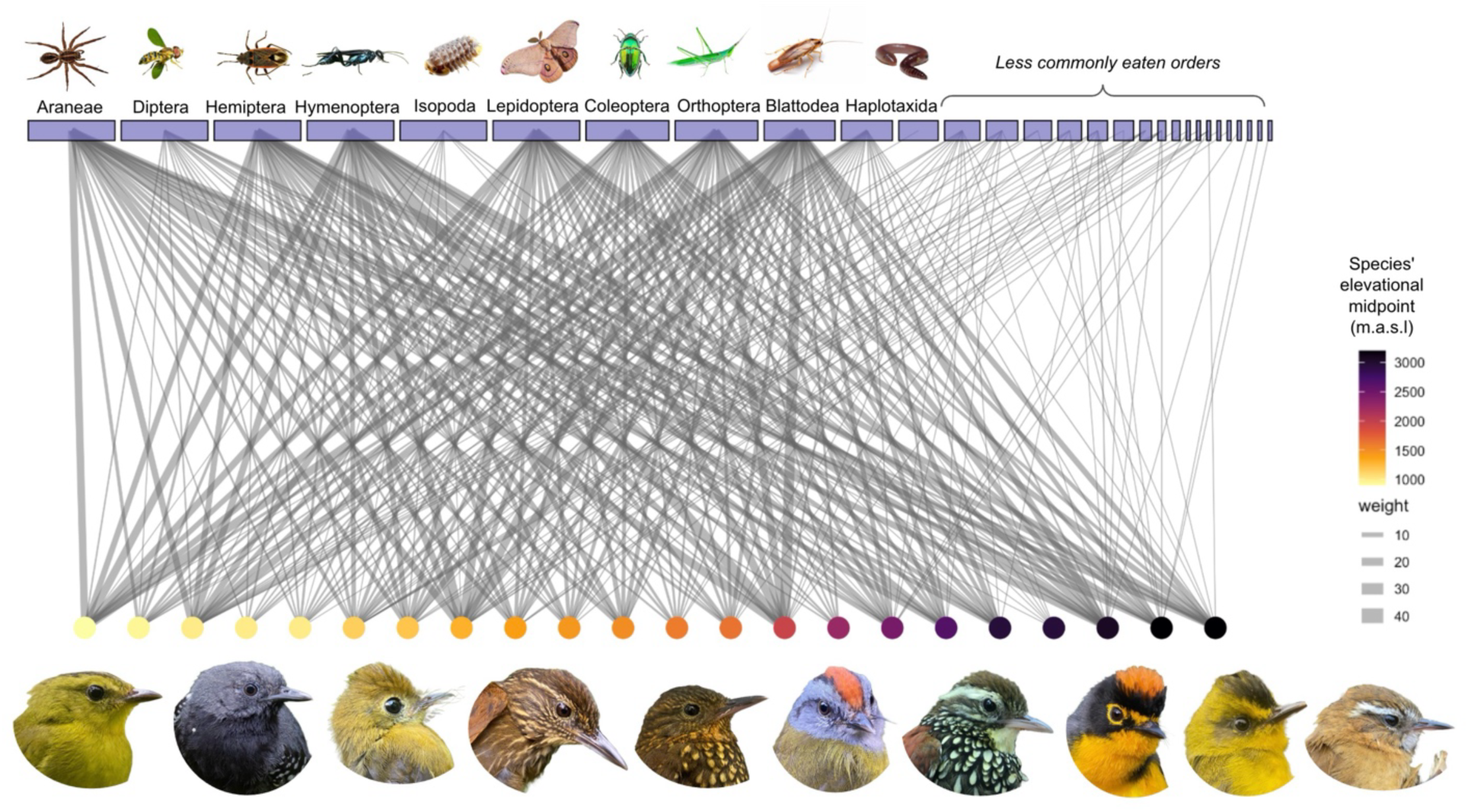
Network of avian predators and their invertebrate prey. Network showing the relative frequency of invertebrate order occurrence in the fecal samples of avian predators. Purple rectangles at the top of the network show prey orders identified by fecal metabarcoding and are sized according to the relative frequency of that prey order across all fecal samples. Colored circles at the bottom of the network correspond to known avian predator species, colored and ordered by the midpoint of the species’ elevational ranges from lowest (light yellow) to highest (dark purple). Gray edges indicate the occurrence of an invertebrate order in a fecal sample collected from the corresponding bird species, weighted by the frequency of occurrence. Bird portraits show a subset of species with representative samples but are also ordered by the midpoint of their elevational range from lowest to highest.

At higher elevations, species were more likely to exhibit probing behavior, flowers and branches were more likely to be used, and Lepidoptera, Diptera, and Chrysomelidae (Coleoptera), and Miridae (Hemiptera) most likely to be consumed. At lower elevations, species were more likely to exhibit gleaning and sallying behavior, attack prey on broad-leafed vegetation, and consume Blattodea and Orthoptera (**Figure 10**). The effect of elevation on the probability of using these resources was similar between our models using a species’ elevational midpoint as a fixed effect and estimating the marginal effect of an individual of a species moving one standard deviation up or down in elevation (**Figure 11**).

**Figure 11:**
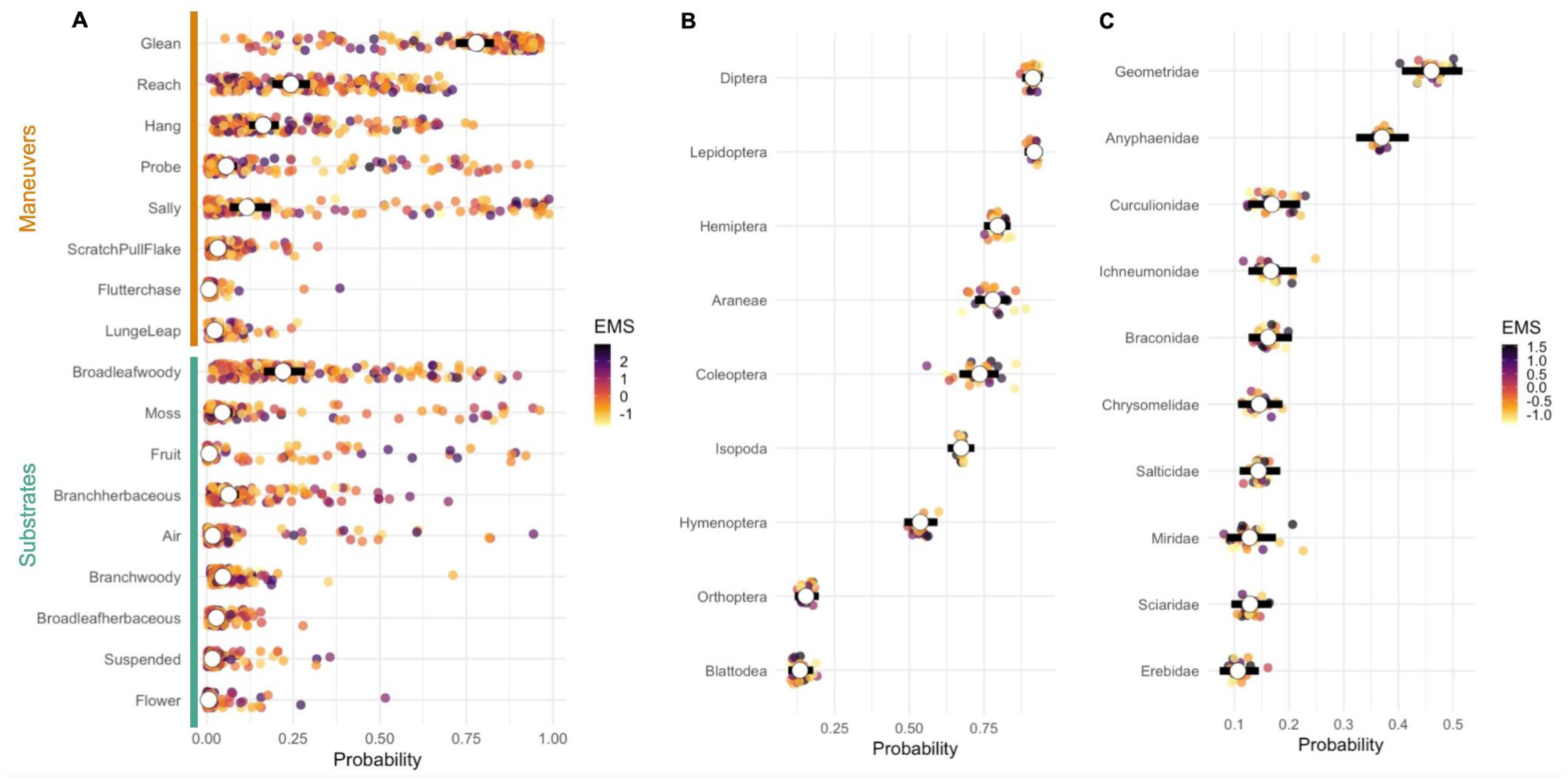
Species vs. community probability intercepts. Estimated probability of (A) performing a foraging maneuver or using a foraging substrate or consuming an invertebrate order (B) or family (C) based on correlated Bernoulli trials in a hierarchical Bayesian model. White dots show the community wide estimated probability, with the black bar showing 95% credible intervals. Colored points show species-specific deviations from the community estimate for each category and are colored by each species’ elevational midpoint. Probabilities are converted from log-odds.

#### Covariation of prey taxa and behaviors

Our correlated Bernoulli models found minimal support for covariation in taxonomic composition within fecal samples (**Figure 12**). However, we found strong support for covariation between many foraging behaviors and substrates within an observation. We found particularly strong correlations between “probing”, “hanging”, “scratching/pulling/flaking”, and moss; and “sally” and air. We also recovered negative covariance between “probing” and “gleaning”, and between broadleaf leaves of woody plants and moss, indicating tradeoffs in a species’ ability to use these behaviors and substrates simultaneously.

**Figure 12:**
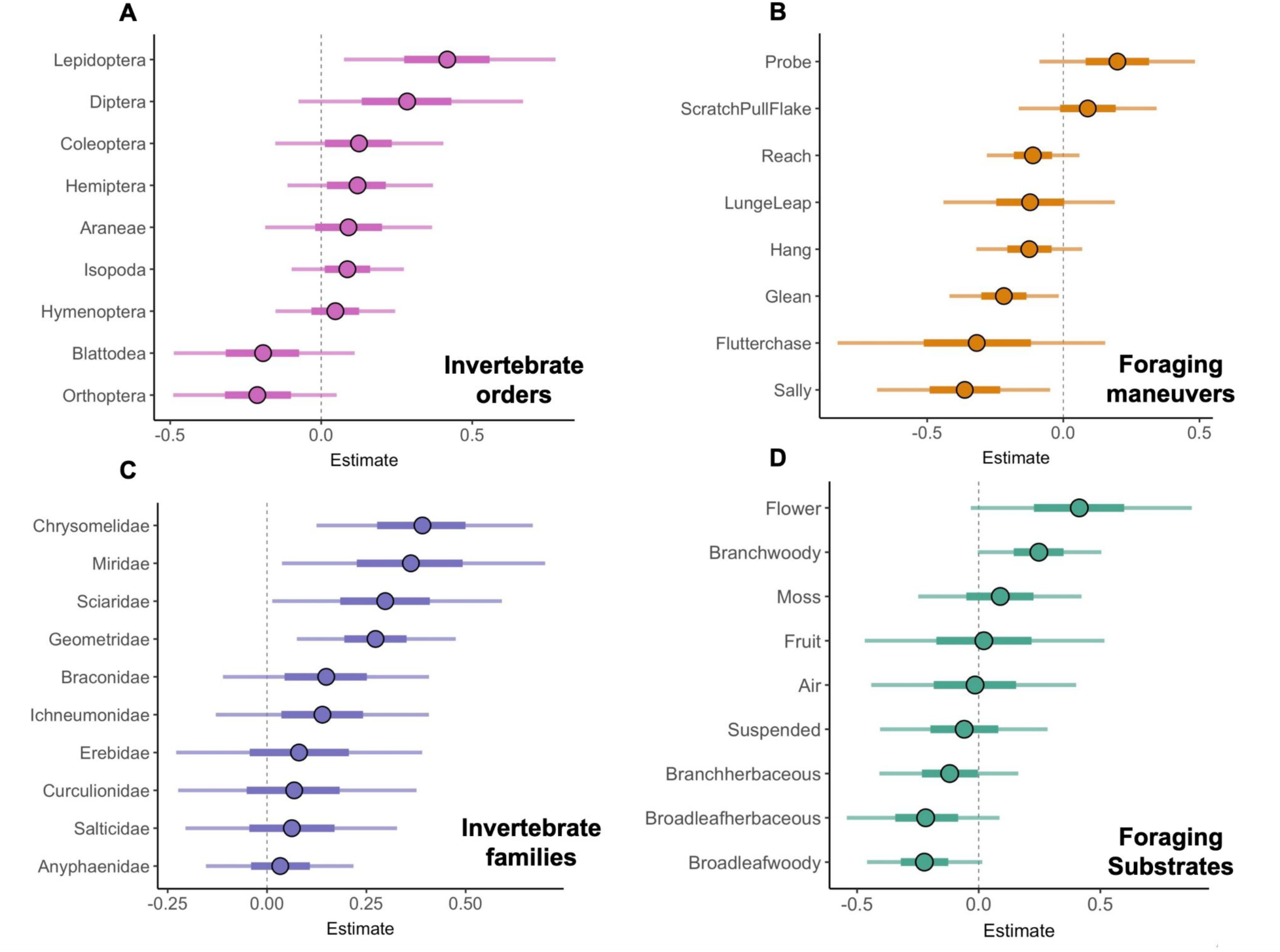
Slope estimates of fixed effect of elevation on diet and foraging behavior. Effect of a species’ elevational midpoint on the probability of performing a foraging behavior (B), using a foraging substrate (D), or consuming an invertebrate order (A) or family (C). Points are mean estimates of the fixed effect slope from correlated Bernoulli trials in a Bayesian mixed effects model, with species and phylogenetic covariance included as random effects (see **Table 2**). Lines show 25-75% and 5-95% credible intervals.

#### Species and Phylogenetic effects

Species identity and phylogenetic similarity captured more than three times as much variation in foraging behavior than prey consumption on average (**Figure 13**, **Table 2**). Specifically, our correlated Bernoulli mixed effect models found that a bird species and phylogenetic similarity underlie approximately 0.3 and 0.4 standard deviations respectively from the community-level probability of consuming an invertebrate order or family. In contrast, deviations from community-level probabilities varied by approximately one full standard deviation based on species, and two standard deviations when species were more closely related. Phylogenetic similarity had a particularly large effect on whether species used “sallying” or “probing” behaviors or “fruit”, “flower”, and “moss” substrates (**Figure 13**).

**Figure 13:**
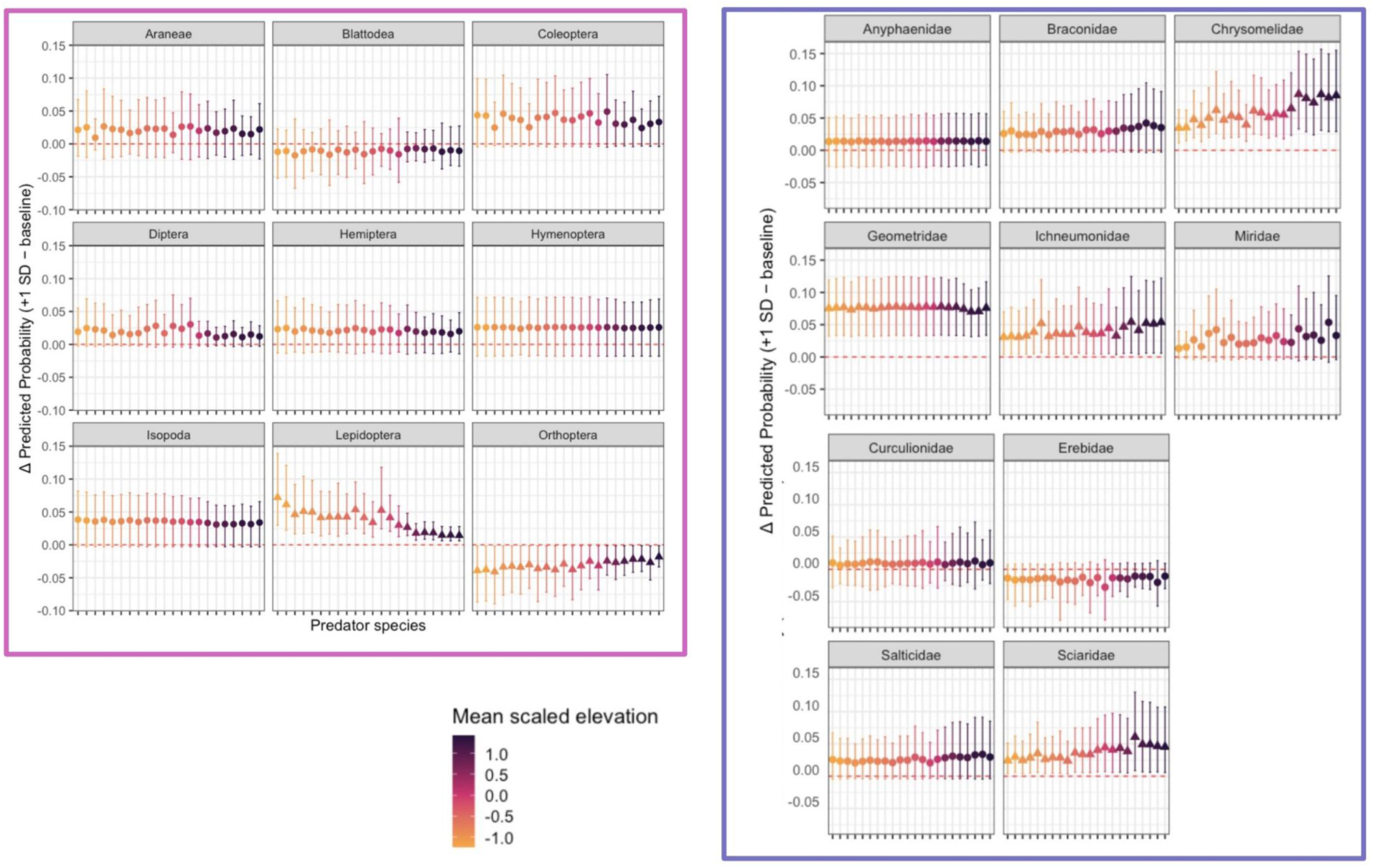
Marginal effects of elevation on intraspecific diet competition. Marginal effect of moving up one standard deviation in elevation on the probability that an individual bird consumed a given invertebrate order (panels boxed in pink) or family (panels boxed in violet). Each point is a bird species, ordered along the x axis by the midpoint of their elevational range from lowest to highest, with 95% credible intervals. Points with credible intervals non-overlapping zero are shaped as triangles.

**Figure 14:**
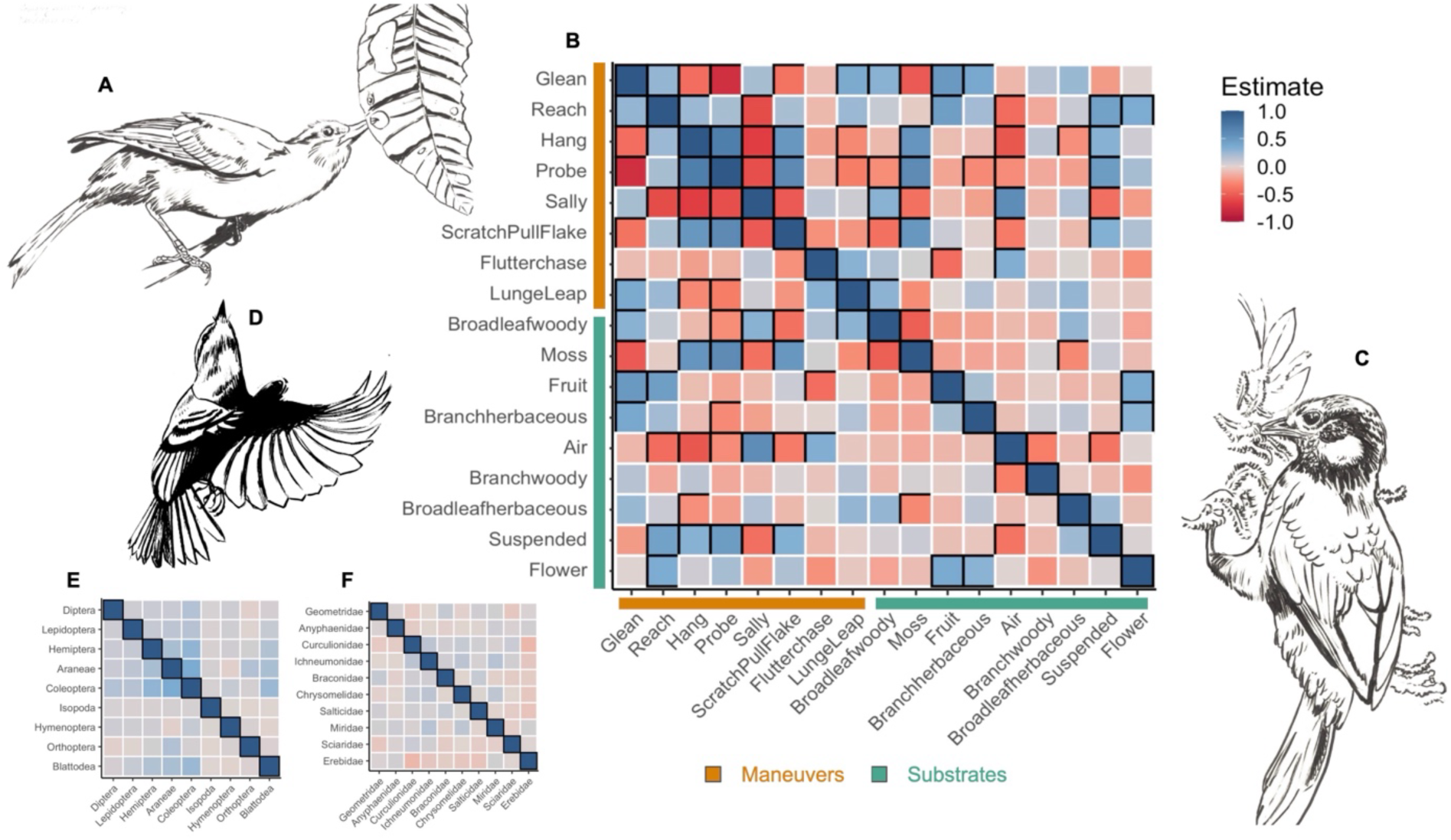
Covariation matrix of foraging behavior and prey selection. Correlation between the co-occurrence of attack maneuvers and substrates within a foraging observation across the Pichincha gradient based on correlated Bernoulli trials (B). Cells bordered with black have 95% credible intervals non-overlapping with zero. Estimate refers to the positive or negative correlation value. Panels E and F show the same parameters between prey orders and families respectively, but we did not detect any significant correlations. Illustrations show key axes of foraging behavior, gleaning (A), probing (C) and sallying (D). Original art by Alexandra Dube.

**Figure 15:**
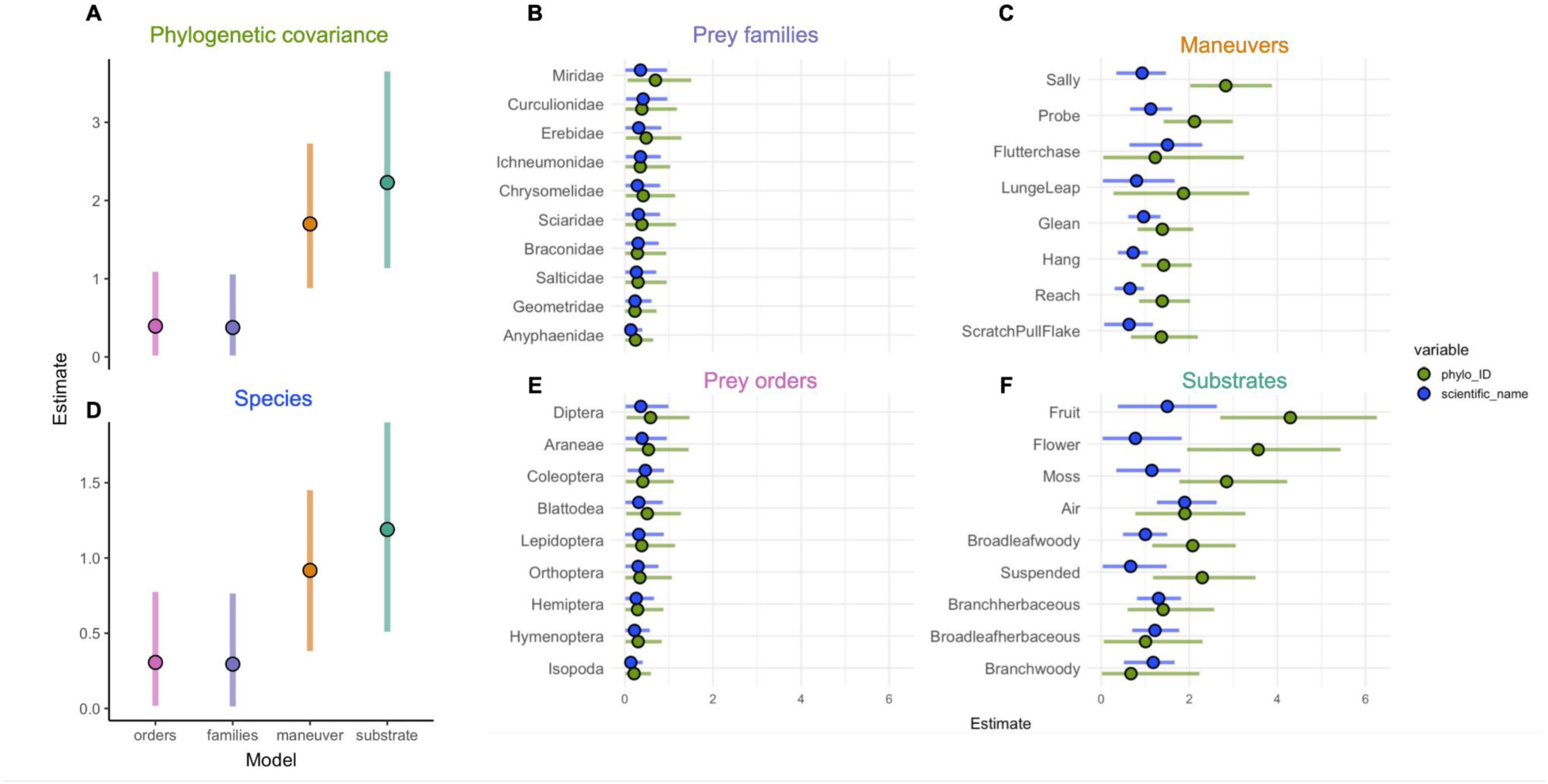
Variance in resource use probabilities induced by species and phylogeny. Variation in resource use probabilities associated with the random effect of species (D) and their phylogenetic similarity (A). Points represent the mean estimate of standard deviations around the community-level intercept for each prey item, foraging maneuver or substrate. Category-specific variation for each model is shown in panels B, C, E and F.

## Discussion

Tropical mountains are the most biodiverse regions on earth because of the stacking of species’ elevational ranges and packing of species into limited niche space (Cadena et al. 2012, Rahbek et al. 2019). Decades of theoretical and empirical work have demonstrated the role of fundamental processes in generating the patterns of ‘stacking’ and ‘packing’, from basic principles of orography to climatic niche evolution and competition (Elsen and Tingley 2015; Cadena et al. 2012; Freeman et al. 2022). However, a mechanistic understanding of these abiotic and biotic processes is limited by the paucity of information about how species acquire resources within the dynamic context tropical mountains (Sherry et al. 2020). Our work addresses this knowledge gap by quantifying *in situ* the dietary and foraging niches of a community of invertivorous birds along an elevational gradient in northwestern Ecuador, finding evidence that community assembly is likely governed more by species-specific behaviors and substrates that birds use to acquire prey than actively selecting particular prey items.

By quantifying resource use directly, our results illuminate the mechanisms of expanding and packing multidimensional niche space across an elevational gradient. Evaluating the volume and density of morphospace in bird communities across quantitatively defined elevational bins recovered the well-documented pattern of broader and more saturated niche space at lower to mid-elevations, particularly in lower cloud forests where lowland and montane communities blend into each other (Pigot et al. 2016; Schumm et al. 2020). The volume and density of ordinated foraging behavior tells a similar story of saturated niches at lower elevations, but with a key difference: species centroids of foraging behavior were further apart in Euclidean space at lower elevations than higher elevations while they were closer together in morphological Euclidean space. That is, species appear to forage more differently from each other at the elevations where they are more morphologically similar, which may facilitate the observed patterns of niche packing. Both the nearest neighbor distances and volume of foraging behavior peak in lower cloudforests, suggesting that a behavioral offset to morphological similarity may promote a wider use of foraging niche space. We also found evidence for dietary niche expansion at lower elevations, supported by the pattern of high-elevation invertebrate prey communities representing a subset of low-elevation communities at the order level (**Figure 4**). This pattern of prey community nestedness across elevational bins in our study system parallels the documented pattern of decreasing variety in invertebrate size and biomass at higher elevations (Guevara and Avilés 2007; Hanz et al. 2019; Sam et al. 2019).

Of all the foraging and diet niche metrics we quantified, conspecific diet similarity had the clearest relationship with elevation. We found a signal of within-species diet composition becoming more similar at high elevations, which is the pattern expected given reduced prey diversity and community composition at higher elevations and matches the recovered pattern of community nestedness (**Figure 4**, **Figure 6**). This pattern was consistent for both invertebrate orders and families, and when considering the effects of species identity and phylogenetic similarity, which explained much less variance in our models of dietary similarity than dietary breadth (**Table 1**). In contrast, phylogenetic similarity of species explained a large amount of variation within-species in foraging behavior, while elevation had a negligible effect, suggesting that prey items are consumed randomly based their availability at a given elevation and that foraging behavior is much more strongly shaped by evolutionary history and morphological constraints.

Despite recovering a general decrease in prey diversity in higher elevational bins, we found that prey diversity within an individual sample generally increases with elevation, suggesting that dietary niche breadth may be more limited by competition within packed niche space at lower elevations than available prey diversity (**Figure 5a**). However, including species and phylogenetic covariance as random effects when modeling the fixed effect of elevation clouded this relationship, weakening the positive effect of elevation on diet breadth at the invertebrate family level and suggesting a negative effect of elevation on diet breadth at the invertebrate order level (**Figure 5b**). That is, variation in diet breadth between species appears to override variation associated with the elevations where a given species lives. Species identity also added noise to the generally negative but weak relationship between elevation and foraging niche breadth (**Figure 5c**). Although subtle, this trend of broader foraging niche breadths at lower elevations may contribute to greater differences in how species forage on average at lower elevations (**Figure 3a**). Taken together, our findings that both within-species foraging niche breadth and nearest neighbor distances between species centroids in ordinated foraging space are greater at lower elevations suggests that morphological niche packing may be facilitated by species diversifying their foraging strategies instead of specializing on few strategies (Klopfer and MacArthur 1961; Sherry et al. 2020). However, disentangling the extent to which these broader foraging niches at lower elevations are a response to within and between species competitive pressures or elevational differences in vegetation structure remains challenging.

Modeling the probability of consuming a particular prey item or using a foraging behavior adds further credence to the hypothesis of that prey is randomly selected based on what is available at a given elevation while evolutionary history and corresponding morphological constraints shape foraging behavior. Species-level estimates of the probability of using a foraging maneuver or substrate often varied dramatically from the community-level estimate, while species varied little around the community-level probability of consuming an invertebrate taxon (**Figure 9**), with species identity and phylogenetic similarity capturing approximately three times as much variation in foraging behavior than prey consumption on average (**Figure 13**). Phylogenetic signal was particularly strong for foraging maneuvers requiring specialized morphology for locomotion or substrate manipulation (**Figure 13c**). Specifically, “sallying” requires long wings for pursuing insects in flight, “lunging or leaping” requires longer legs for sufficient speed and leverage, and “probing” often requires a longer bill for reaching inside substrates. While wing, tarsus, and bill length are all relatively malleable over shallow and deep time, phylogenetic conservatism of these traits has a clear effect on how birds access prey (Sayol et al. 2025). Phylogenetic signal was similarly strong for several foraging substrates, likely reflecting both the association with the maneuvers birds use to interface with the substrate and taxa-specific preferences (**Figure 13f**). For example, moss typically requires manipulation with beaks or feet, while flowers were more frequently used at higher elevations, primarily by species in the omnivorous family Thraupidae that is common at higher elevations (Naoki 2007; Burns et al. 2014). We can further disentangle the relationship between morphology and substrate use based on the correlation estimates from the Bernoulli models, which highlight how many maneuvers and substrates may be mutually exclusive (**Figure 12**). Among the substrates used by more closely related species, moss and ‘probing’ were tightly correlated but flowers and fruit were not strongly associated with phylogenetically conserved maneuvers, suggesting a morphological constraint on using some, but not all foraging substrates.

Constraints on foraging strategies also highlight cases where elevation does seem to affect resource use. As a key example, ‘gleaning’ and ‘probing’ were mutually exclusive within a single foraging observation (**Figure 12**). While the tradeoff between these behaviors is intuitive (a bird cannot attack a prey item on the surface of a substrate and inside the substrate at the same time), we found that gleaning is significantly more common at lower elevations while probing is more common at higher elevations (**Figure 10b**). The substrates associated with these maneuvers, primarily the leaves of woody plants with “gleaning” and moss with “probing”, showed the corresponding negative and positive relationship with elevation respectively (**Figure 10d**). In addition to morphological differences between species, this glean-probe tradeoff and other elevational shifts in maneuver and substrate use may be driven by changes in forest structural complexity with elevation (Remsen Jr 1985; Davies et al. 2007; Ferger, Schleuning, Hemp, Howell, and Böhning-Gaese 2014). The reduction in canopy height, leaf size, and herbaceous vegetation at higher elevations is a well-documented pattern that we also found in our system by visually estimating the canopy height and size of the leaves where birds foraged (Jankowski, Merkord, et al. 2013, Figure S5). Less broad-leaf substrate and vertical strata to partition at higher elevations may promote the use of other substrates such as flowers, branches, and to some extent moss at higher elevations (**Figure 10d**). This shift in habitat structure at higher elevations may also underlie our finding of a subtle reduction in foraging niche breadths of higher-elevation species (**Figure 4c**).

Elevation also influenced the probability of consuming several prey taxa (**Figure 10a,c**), but the lack of both correlated prey consumption within samples and phylogenetic signal in predicting their occurrence in a fecal sample suggests that birds are more likely to be randomly eating available prey rather than selecting for specific taxa. Within-species patterns support this hypothesis, with the probability of a bird species consuming a prey taxon either not changing with elevation or paralleling between-species patterns (**Figure 11**). That is, the same bird species appear equally likely to eat the same items at different elevations, except for prey taxa that are locally more common at specific elevations. This apparent indifference of birds to specific prey taxa complements other studies that have quantified the biomass and body size of arthropods along elevational gradients regardless of their taxonomic identity, finding a negative relationship between arthropod abundance and size with elevation and the species richness and functional diversity of birds (Sam et al. 2019, Hanz et al. 2019, Schumm et al. 2020). Considered alongside these studies, our work strongly implies that shifts in invertivorous bird abundance and distribution along elevational gradients are not in pursuit of particular prey taxa, but may be responses to a general lack of food and other resources associated with climate or land use change (Ferger 2014; Freeman et al. 2018; Neate-Clegg et al. 2021; Newell et al. 2023)

While directly sampling invertebrate diversity, abundance, and size at our field sites was outside the scope of this study, our findings nonetheless capture several known elevational patterns in Arthropod diversity. The generally large-bodied orders Blattodea and Orthoptera were more likely to be consumed at lower elevations, matching expectations of bigger arthropods at lower elevations (Supriya et al. 2019), **Figure 10a**). We also capture the prevalence of Diptera at higher elevations, as did a classic study of arthropod diversity patterns elsewhere in the Andes (Janzen et al. 1976). However, our finding that leaf-beetles in the family Chrysomelidae were most likely to be consumed at higher elevations contradicts a study of this family from the eastern slope of the Ecuadorian Andes, which found them to be more common and diverse at lower elevations (Thormann et al. 2018). Furthermore, the strongest caveat to our dataset derived from DNA metabarcoding is that it is naive to the life stage of the prey item (e.g. caterpillar vs adult butterfly) and thus offers reduced insight into the relationship between prey, foraging behavior, and functional morphology (Hoenig et al. 2022). For example, our data cannot resolve the extent to which the most consumed invertebrate family, Geometrid moths, were captured in their adult or larval forms. In some cases, functional traits may strongly indicate which form was taken. For example, if a species is more likely to probe in moss than sally from the air then it is arguably more likely to capture a flying insect in larval than adult form, but this is not directly testable with our dataset. Differences in life stages may also affect biases in metabarcoding, with soft-bodied larval forms more likely to have higher read counts than chitinous adult forms. Despite these challenges, the scalability of DNA metabarcoding across many sites and taxa make it an invaluable tool for quantifying dietary niches.

By integrating field observations of foraging behavior with diet information revealed from DNA metabarcoding, we demonstrate how foraging behavior facilitates the packing of morphologically similar species into niche space along the slopes of the tropical Andes, contributing to extreme species richness. Our findings also suggest that foraging behavior, not prey identity, is the dimension of resource use that varies most between species and likely drives key processes of community assembly such as competition, despite detectable shifts in prey community composition, vegetation structure, and competition intensity across the elevational gradient in our study system. As ecological communities in the tropical Andes and other global biodiversity hotspot restructure in the face of climate and land use change, our findings urge the conservation of forests with sufficient habitat structure that can support diverse assemblages of birds and their invertebrate prey.

## Acknowledgements

We are eternally indebted to the plethora of hosts and assistants at field sites, including Fundacion Jocotoco, Nicole Buttner, Dusti Becker, Matteo Roldan, Chiara Correa, Daniela Franco, Guillermo Larco, Roberto Lima, Reserva Intillacta, Nelson Apolo, Juan Carlos Crespo, Leo and Christian Montalvo, and Richard Parsons. We thank our world class field technicians and ornithologists William Arteaga, Gabriela Mena and Maria Jose Arias. We are grateful for Kevin Feldheim in the Pritzker DNA Lab at the Field Museum. We thank Trevor Price and Tim Wootton at the University of Chicago for thoughtful commentary on this manuscript throughout its development. This research was funded by the Tinker Foundation’s Center for Latin American Studies, the American Ornithological Society, the Animal Behavior Society, the American Philosophical Society, the American Museum of Natural History, the Field Museum of Natural History, and the University of Chicago.

---

**Table S1:**
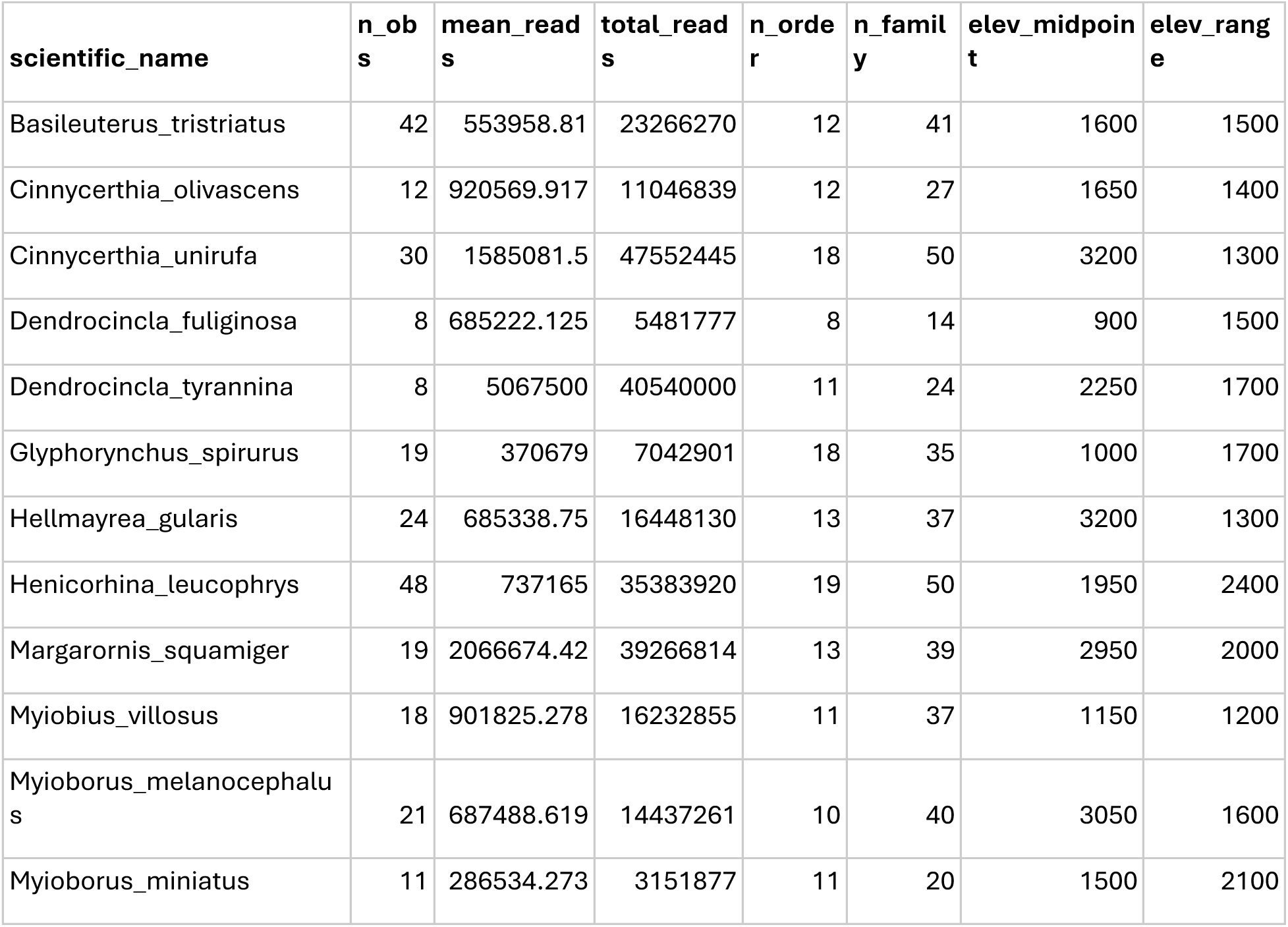
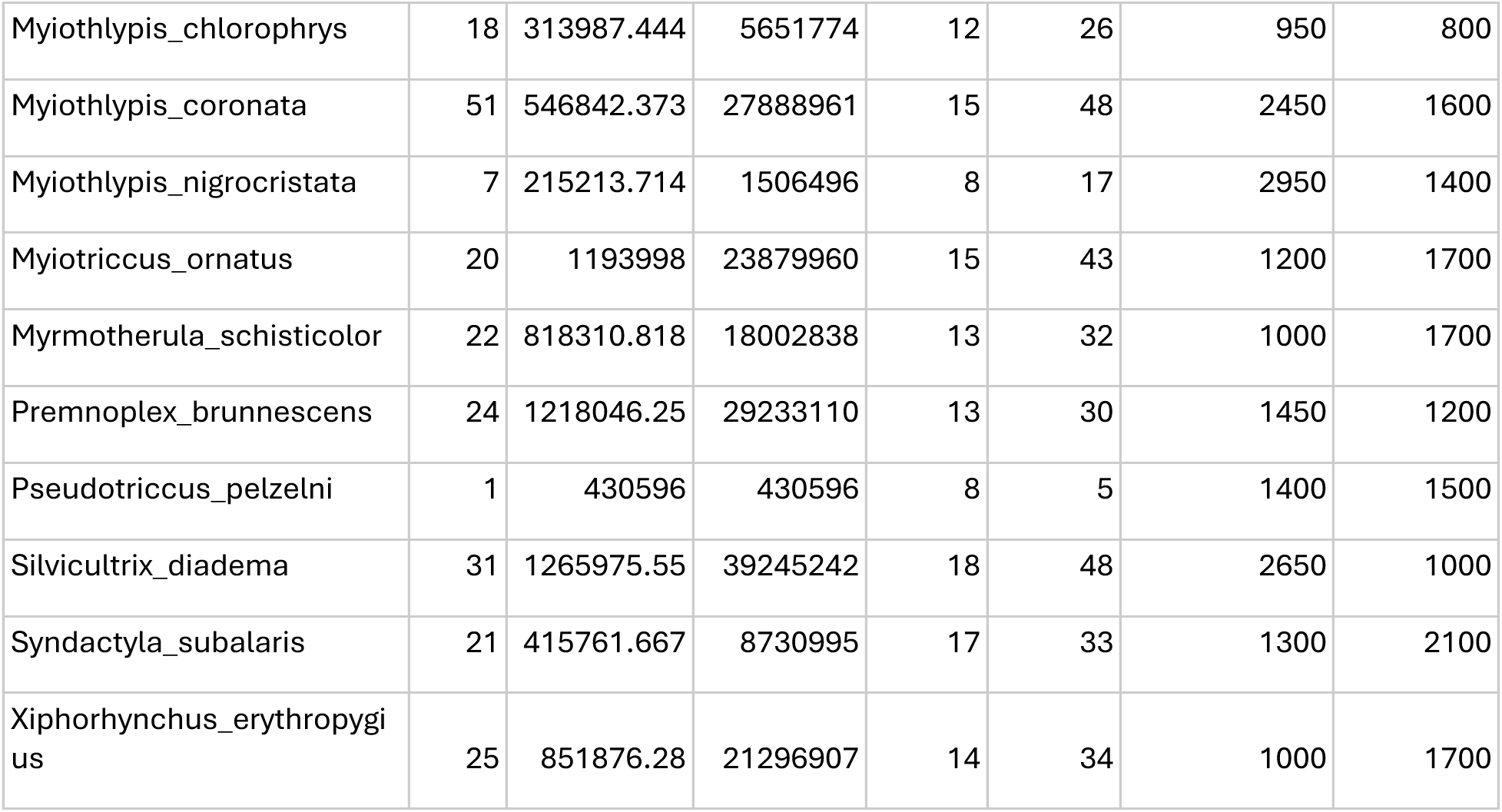
Summary of species with quantified diet.

**Table S2:**
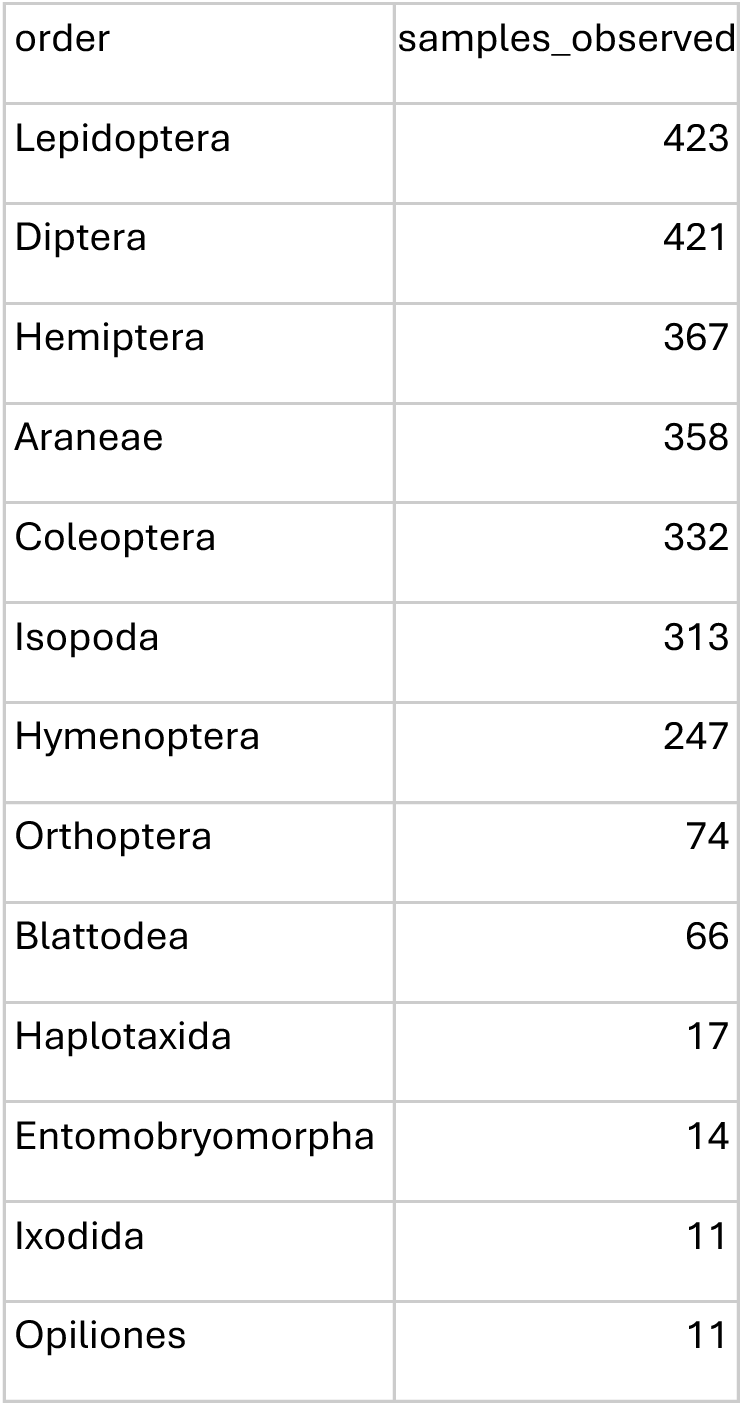
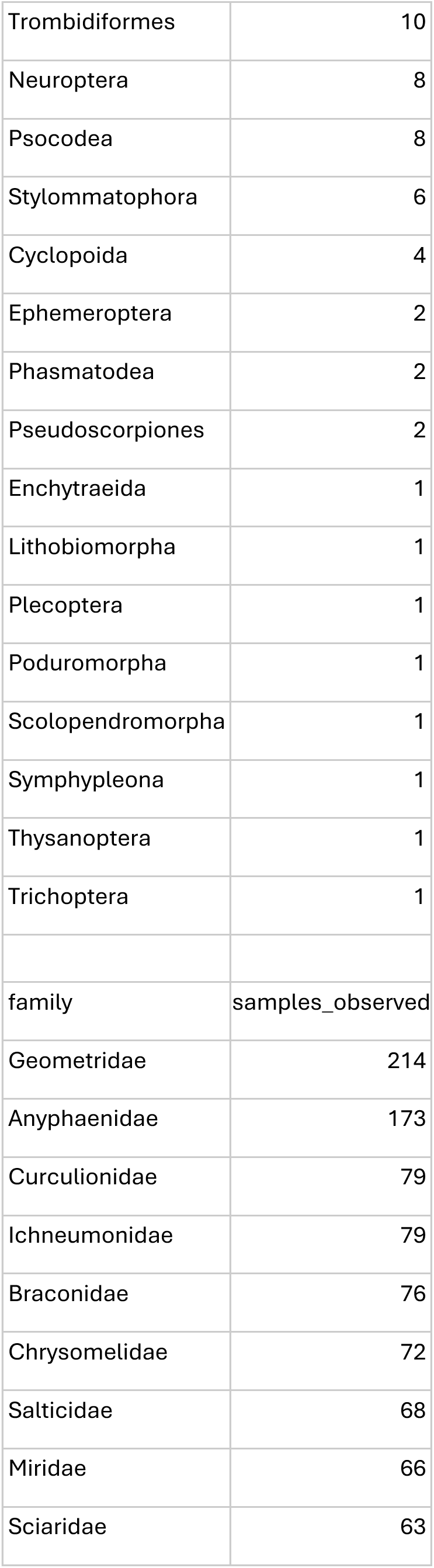
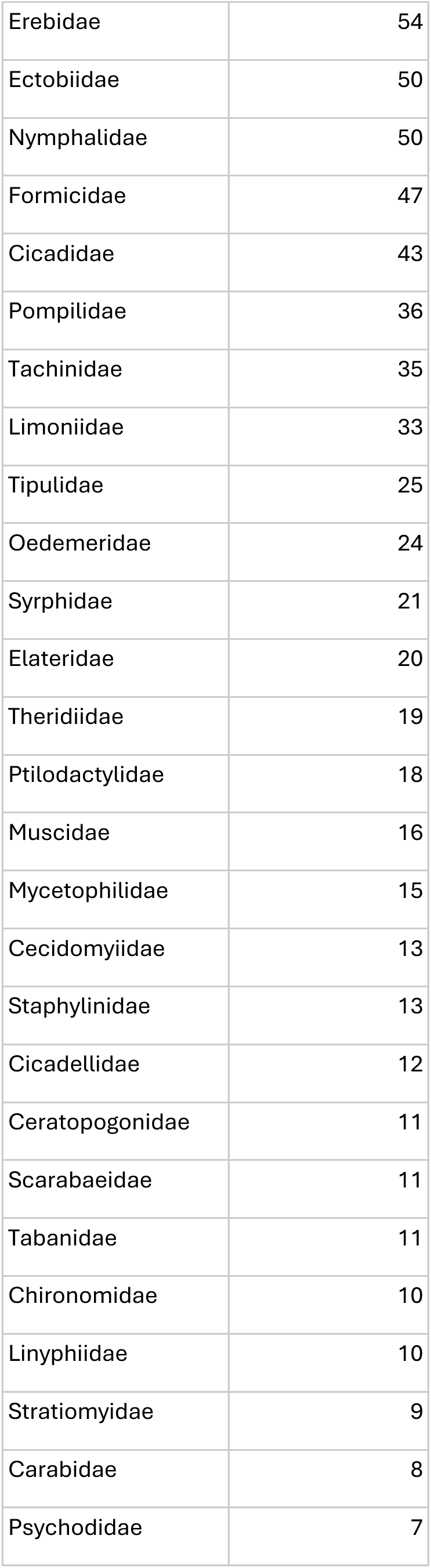
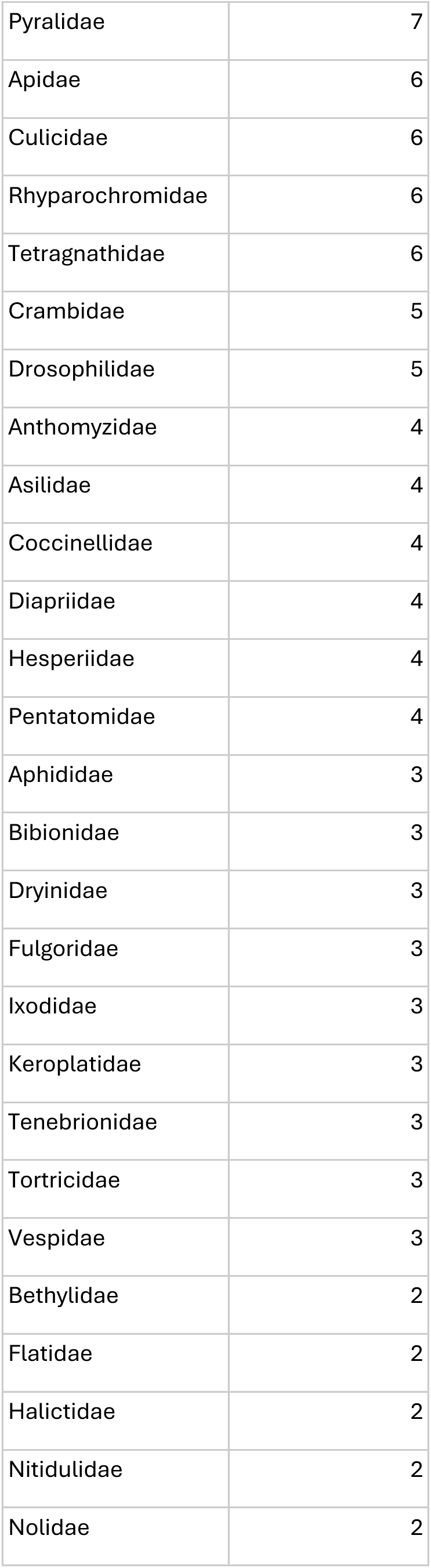
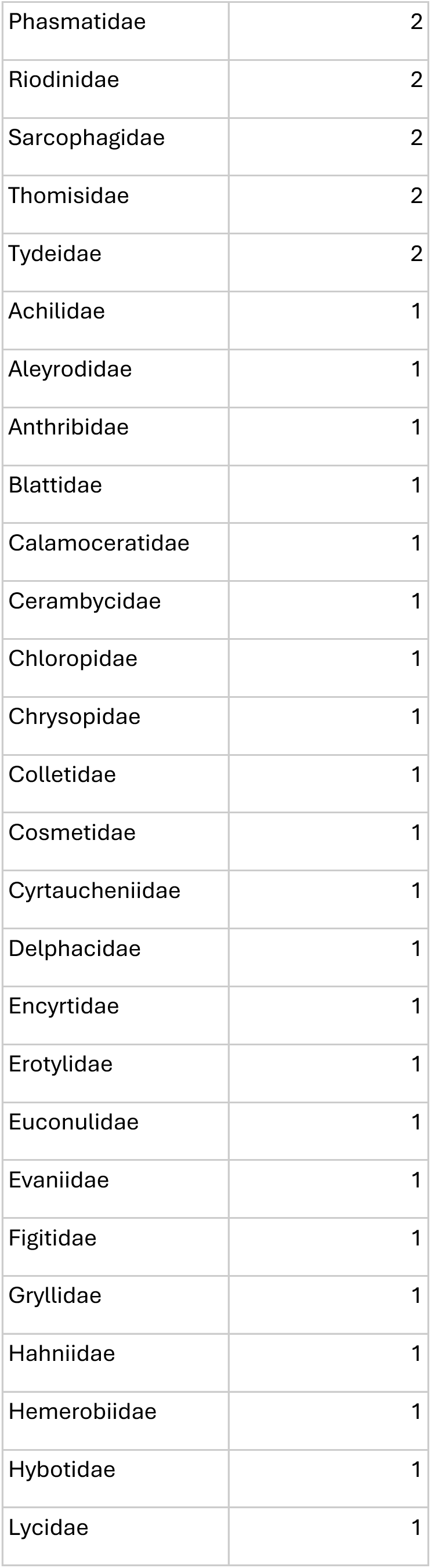
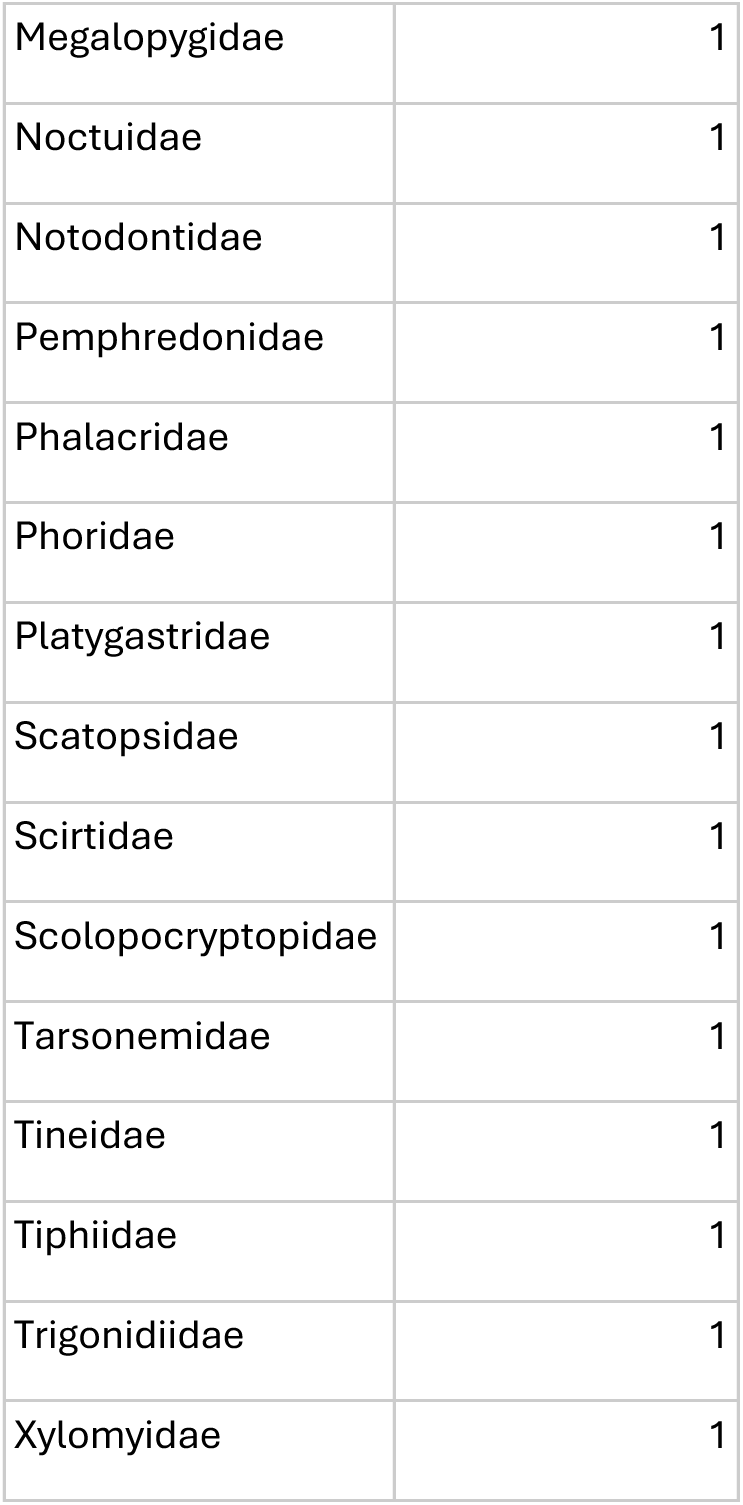
Counts of detected invertebrate prey orders and families.

## Works Cited

Ausprey, Ian J., Felicity L. Newell, and Scott K. Robinson. 2022. ‘Functional Response Traits and Altered Ecological Niches Drive the Disassembly of Cloud Forest Bird Communities in Tropical Montane Countrysides’. Journal of Animal Ecology 91 (11): 2314–28. 10.1111/1365-2656.13816.

Bennett, Joanne M., Jennifer Sunday, Piero Calosi, et al. 2021. ‘The Evolution of Critical Thermal Limits of Life on Earth’. Nature Communications 12 (1): 1. 10.1038/s41467-021-21263-8.

Blonder, Benjamin, Cecina Babich Morrow, Brian Maitner, et al. 2018. ‘New Approaches for Delineating N-Dimensional Hypervolumes’. Methods in Ecology and Evolution 9 (2): 305–19. 10.1111/2041-210X.12865.

Bonetti, Maria Fernanda, and John J. Wiens. 2014. ‘Evolution of Climatic Niche Specialization: A Phylogenetic Analysis in Amphibians’. Proceedings of the Royal Society B: Biological Sciences 281 (1795): 20133229.

Bürkner, Paul-Christian. 2017. ‘Brms: An R Package for Bayesian Multilevel Models Using Stan’. Journal of Statistical Software 80 (August): 1–28. 10.18637/jss.v080.i01.

Burns, Kevin J., Allison J. Shultz, Pascal O. Title, et al. 2014. ‘Phylogenetics and Diversification of Tanagers (Passeriformes: Thraupidae), the Largest Radiation of Neotropical Songbirds’. Molecular Phylogenetics and Evolution 75 (June): 41–77. 10.1016/j.ympev.2014.02.006.

Cadena, Carlos Daniel, and Laura N. Céspedes. 2020. ‘Origin of Elevational Replacements in a Clade of Nearly Flightless Birds: Most Diversity in Tropical Mountains Accumulates via Secondary Contact Following Allopatric Speciation’. In Neotropical Diversification: Patterns and Processes. Springer.

Cadena, Carlos Daniel, Kenneth H. Kozak, Juan Pablo Gómez, et al. 2012. ‘Latitude, Elevational Climatic Zonation and Speciation in New World Vertebrates’. Proceedings of the Royal Society B: Biological Sciences 279 (1726): 194–201. 10.1098/rspb.2011.0720.

Cavender-Bares, Jeannine, Antonio Gonzalez-Rodriguez, Annette Pahlich, Kari Koehler, and Nicholas Deacon. 2011. ‘Phylogeography and Climatic Niche Evolution in Live Oaks (Quercus Series Virentes) from the Tropics to the Temperate Zone’. Journal of Biogeography 38 (5): 962–81.

Colwell, Robert K., and Kenneth J. Feeley. 2025. ‘Still Little Evidence of Poleward Range Shifts in the Tropics, but Lowland Biotic Attrition May Be Underway’. Biotropica 57 (1): e13358. 10.1111/btp.13358.

Crouch, Nicholas M. A., João M. G. Capurucho, Shannon J. Hackett, and John M. Bates. 2019. ‘Evaluating the Contribution of Dispersal to Community Structure in Neotropical Passerine Birds’. Ecography 42 (2): 390–99. 10.1111/ecog.03927.

Davies, Richard G, C. David L Orme, David Storch, et al. 2007. ‘Topography, Energy and the Global Distribution of Bird Species Richness’. Proceedings of the Royal Society B: Biological Sciences 274 (1614): 1189–97. 10.1098/rspb.2006.0061.

Dey, Kushal K., Chiaowen Joyce Hsiao, and Matthew Stephens. 2017. ‘Visualizing the Structure of RNA-Seq Expression Data Using Grade of Membership Models’. PLOS Genetics 13 (3): e1006599. 10.1371/journal.pgen.1006599.

Dixon, Philip. 2003. ‘VEGAN, a Package of R Functions for Community Ecology’. Journal of Vegetation Science 14 (6): 927–30. 10.1111/j.1651103.2003.tb02228.x.

Dobzhansky, Theodosius. 1950. ‘Evolution in the Tropics’. Am. Sci. 38: 209–21.

Elsen, Paul R., and Morgan W. Tingley. 2015. ‘Global Mountain Topography and the Fate of Montane Species under Climate Change’. Nature Climate Change 5 (8): 772–76.

Endler, John A. 1982. ‘Problems in Distinguishing Historical from Ecological Factors in Biogeography’. American Zoologist 22 (2): 441–52.

Ferger, Stefan W., Matthias Schleuning, Andreas Hemp, Kim M. Howell, and Katrin Böhning-Gaese. 2014. ‘Food Resources and Vegetation Structure Mediate Climatic Effects on Species Richness of Birds’. Global Ecology and Biogeography 23 (5): 541–49. 10.1111/geb.12151.

Ferger, Stefan W., Matthias Schleuning, Andreas Hemp, Kim M. Howell, and Katrin Böhning-Gaese. 2014. ‘Food Resources and Vegetation Structure Mediate Climatic Effects on Species Richness of Birds’. Global Ecology and Biogeography 23 (5): 541–49.

Forsman, Anna M, Brandon D Hoenig, Stephanie A Gaspar, Jason D Fischer, Joe Siegrist, and Kevin Fraser. 2022. ‘Evaluating the Impacts of Metabarcoding Primer Selection on DNA Characterization of Diet in an Aerial Insectivore, the Purple Martin’. Ornithology 139 (1): ukab075. 10.1093/ornithology/ukab075.

Freeman, Benjamin G. 2015. ‘Competitive Interactions upon Secondary Contact Drive Elevational Divergence in Tropical Birds’. The American Naturalist 186 (4): 470–79. 10.1086/682703.

Freeman, Benjamin G., Micah N. Scholer, Viviana Ruiz-Gutierrez, and John W. Fitzpatrick. 2018. ‘Climate Change Causes Upslope Shifts and Mountaintop Extirpations in a Tropical Bird Community’. Proceedings of the National Academy of Sciences 115 (47): 11982–87. 10.1073/pnas.1804224115.

Freeman, Benjamin G., Yiluan Song, Kenneth J. Feeley, and Kai Zhu. 2021. ‘Montane Species Track Rising Temperatures Better in the Tropics than in the Temperate Zone’. Ecology Letters 24 (8): 1697–708. 10.1111/ele.13762.

Freeman, Benjamin G., Matthew Strimas-Mackey, and Eliot T. Miller. 2022. ‘Interspecific Competition Limits Bird Species’ Ranges in Tropical Mountains’. Science 377 (6604): 416–20. 10.1126/science.abl7242.

García-Moreno, Jaime, Peter Arctander, and Jon Fjeldsaa. 1999. ‘Strong Diversification at the Treeline among Metallura Hummingbirds’. The Auk 116 (3): 702–11.

Ghosh-Harihar, Mousumi, and Trevor D. Price. 2014. ‘A Test for Community Saturation along the Himalayan Bird Diversity Gradient, Based on within-Species Geographical Variation’. Journal of Animal Ecology, 628–38.

Glenn, Travis C., Todd W. Pierson, Natalia J. Bayona-Vásquez, et al. 2019. ‘Adapterama II: Universal Amplicon Sequencing on Illumina Platforms (TaggiMatrix)’. PeerJ 7 (October): e7786. 10.7717/peerj.7786.

Guevara, Jennifer, and Leticia Avilés. 2007. ‘Multiple Techniques Confirm Elevational Differences in Insect Size That May Influence Spider Sociality’. Ecology 88 (8): 2015–23.

Hanz, Dagmar, Katrin Böhning-Gaese, Stefan Ferger, et al. 2019. ‘Functional and Phylogenetic Diversity of Bird Assemblages Are Filtered by Different Biotic Factors on Tropical Mountains’. Journal of Biogeography 46 (February): 291–303. 10.1111/jbi.13489.

Harvey, Michael G., Gustavo A. Bravo, Santiago Claramunt, et al. 2020. ‘The Evolution of a Tropical Biodiversity Hotspot’. Science 370 (6522): 1343–48. 10.1126/science.aaz6970.

Hennig, Sabine, Robert Vogler, and Jiří Pánek. 2023. ‘Survey123 for ArcGIS Online’. In Evaluating Participatory Mapping Software, edited by Charla M. Burnett. Springer International Publishing. 10.1007/978-3-031-19595_8.

Hoenig, Brandon D, Allison M Snider, Anna M Forsman, et al. 2022. ‘Current Methods and Future Directions in Avian Diet Analysis’. Ornithology 139 (1): ukab077. 10.1093/ornithology/ukab077.

Hua, Xia. 2016. ‘The Impact of Seasonality on Niche Breadth, Distribution Range and Species Richness: A Theoretical Exploration of Janzen’s Hypothesis’. Proceedings of the Royal Society B: Biological Sciences 283 (1835): 20160349.

Huey, Raymond B. 1978. ‘Latitudinal Pattern of Between-Altitude Faunal Similarity: Mountains Might Be” Higher” in the Tropics’. The American Naturalist 112 (983): 225–29.

Jankowski, Jill E., Gustavo A. Londoño, Scott K. Robinson, and Mark A. Chappell. 2013. ‘Exploring the Role of Physiology and Biotic Interactions in Determining Elevational Ranges of Tropical Animals’. Ecography 36 (1): 1–12. 10.1111/j.1600-0587.2012.07785.x.

Jankowski, Jill E., Christopher L. Merkord, William Farfan Rios, Karina García Cabrera, Norma Salinas Revilla, and Miles R. Silman. 2013. ‘The Relationship of Tropical Bird Communities to Tree Species Composition and Vegetation Structure along an Andean Elevational Gradient’. Journal of Biogeography 40 (5): 950–62. 10.1111/jbi.12041.

Janzen, D. H., M. Ataroff, M. Farinas, et al. 1976. ‘Changes in the Arthropod Community along an Elevational Transect in the Venezuelan Andes’. Biotropica 8 (3): 193. 10.2307/2989685.

Janzen, Daniel H. 1967. ‘Why Mountain Passes Are Higher in the Tropics’. The American Naturalist 101 (919): 233–49.

Jetz, Walter, Holger Kreft, Gerardo Ceballos, and Jens Mutke. 2009. ‘Global Associations between Terrestrial Producer and Vertebrate Consumer Diversity’. Proceedings of the Royal Society B: Biological Sciences 276 (1655): 269–78. 10.1098/rspb.2008.1005.

Klopfer, Peter H., and R. H. MacArthur. 1961. ‘On the Causes of Tropical Species Diversity: Niche Overlap’. The American Naturalist 95 (883): 223–26. 10.1086/282179.

Korejs, Kryštof, Bonny Koane, Samuel Jeppy, Leonardo Ré Jorge, Vojtěch Novotný, and Kateřina Sam. 2025. ‘Feeding Specialisation Shapes Avian Functional Diversity Along a Tropical Rainforest Elevational Gradient’. Journal of Biogeography 52 (5): e15103. 10.1111/jbi.15103.

Laurance, William F., D. Carolina Useche, Luke P. Shoo, et al. 2011. ‘Global Warming, Elevational Ranges and the Vulnerability of Tropical Biota’. Biological Conservation 144 (1): 548–57. 10.1016/j.biocon.2010.10.010.

Linck, Ethan B., Benjamin G. Freeman, C. Daniel Cadena, and Cameron K. Ghalambor. 2021. ‘Evolutionary Conservatism Will Limit Responses to Climate Change in the Tropics’. Biology Letters 17 (10): 20210363. 10.1098/rsbl.2021.0363.

MacArthur, Robert H. 1984. Geographical Ecology: Patterns in the Distribution of Species. Princeton University Press.

MacArthur, Robert, and Richard Levins. 1967. ‘The Limiting Similarity, Convergence, and Divergence of Coexisting Species’. The American Naturalist 101 (921): 377–85.

Marra, Peter P., and J. V. Remsen Jr. 1997. ‘Insights into the Maintenance of High Species Diversity in the Neotropics: Habitat Selection and Foraging Behavior in Understory Birds of Tropical and Temperate Forests’. Ornithological Monographs, 445–83.

May, Robert M., and Robert H. MacArthur. 1972. ‘Niche Overlap as a Function of Environmental Variability’. Proceedings of the National Academy of Sciences 69 (5): 1109–13. 10.1073/pnas.69.5.1109.

McCain, Christy M. 2009. ‘Vertebrate Range Sizes Indicate That Mountains May Be “Higher” in the Tropics’. Ecology Letters 12 (6): 550–60. 10.1111/j.1461-0248.2009.01308.x.

McTavish, Emily Jane, Jeff A. Gerbracht, Mark T. Holder, et al. 2025. ‘A Complete and Dynamic Tree of Birds’. Proceedings of the National Academy of Sciences 122 (18): e2409658122. World. 10.1073/pnas.2409658122.

Moles, Angela T., and Jeff Ollerton. 2016. ‘Is the Notion That Species Interactions Are Stronger and More Specialized in the Tropics a Zombie Idea?’ Biotropica 48 (2): 141–45.

Montaño-Centellas, Flavia A., Bette A. Loiselle, and Morgan W. Tingley. 2020. ‘Ecological Drivers of Avian Community Assembly along a Tropical Elevation Gradient’. Ecography, December 21, ecog.05379. 10.1111/ecog.05379.

Naoki, Kazuya. 2007. ‘Arthropod Resource Partitioning Among Omnivorous Tanagers (Tangara SPP.) in Western Ecuador’. The Auk 124 (1): 197–209. 10.1093/auk/124.1.197.

Neate-Clegg, Montague H, Samuel E Jones, Joseph A Tobias, William D Newmark, and H Şekercioğlu. 2021. Ecological Correlates of Elevational Range Shifts in Tropical Birds. 36.

Newell, Felicity L., Ian J. Ausprey, and Scott K. Robinson. 2023. ‘Wet and Dry Extremes Reduce Arthropod Biomass Independently of Leaf Phenology in the Wet Tropics’. Global Change Biology 29 (2): 308–23. 10.1111/gcb.16379.

O’Rourke, Devon R., Nicholas A. Bokulich, Michelle A. Jusino, Matthew D. MacManes, and Jeffrey T. Foster. 2020. ‘A Total Crapshoot? Evaluating Bioinformatic Decisions in Animal Diet Metabarcoding Analyses’. Ecology and Evolution 10 (18): 9721–39. 10.1002/ece3.6594.

Paradis, Emmanuel, Julien Claude, and Korbinian Strimmer. 2004. ‘APE: Analyses of Phylogenetics and Evolution in R Language’. Bioinformatics 20 (2): 289–90. 10.1093/bioinformatics/btg412.

Pellissier, Vincent, Jean-Yves Barnagaud, W. Daniel Kissling, Çağan Şekercioğlu, and Jens-Christian Svenning. 2018. ‘Niche Packing and Expansion Account for Species Richness–Productivity Relationships in Global Bird Assemblages’. Global Ecology and Biogeography 27 (5): 604–15. 10.1111/geb.12723.

Pigot, Alex L., and Joseph A. Tobias. 2015. ‘Dispersal and the Transition to Sympatry in Vertebrates’. Proceedings of the Royal Society B: Biological Sciences 282 (1799): 20141929. 10.1098/rspb.2014.1929.

Pigot, Alex L., Christopher H. Trisos, and Joseph A. Tobias. 2016. ‘Functional Traits Reveal the Expansion and Packing of Ecological Niche Space Underlying an Elevational Diversity Gradient in Passerine Birds’. Proceedings of the Royal Society B: Biological Sciences 283 (1822): 20152013. 10.1098/rspb.2015.2013.

Price, Trevor D., Daniel M. Hooper, Caitlyn D. Buchanan, et al. 2014. ‘Niche Filling Slows the Diversification of Himalayan Songbirds’. Nature 509 (7499): 7499. 10.1038/nature13272.

Rahbek, Carsten, Michael K. Borregaard, Robert K. Colwell, et al. 2019. ‘Humboldt’s Enigma: What Causes Global Patterns of Mountain Biodiversity?’ Science 365 (6458): 1108–13. 10.1126/science.aax0149.

Remsen, J. V., and Scott K. Robinson. 1990. ‘A Classification Scheme for Foraging Behavior of Birds in Terrestrial Habitats’. Studies in Avian Biology 13: 144–60.

Remsen Jr, J. V. 1985. ‘Community Organization and Ecology of Birds of High Elevation Humid Forest of the Bolivian Andes’. Ornithological Monographs, 733–56.

Rosenberg, Gary H. 1990. ‘Habitat Specialization and Foraging Behavior by Birds of Amazonian River Islands in Northeastern Peru’. The Condor 92 (2): 427–43.

Rosenberg, Kenneth V. 1990. ‘Dead-Leaf Foraging Specialization in Tropical Forest Birds: Measuring Resources Availability and Use’. Studies in Avian Biology 13: 360–68.

Sam, Katerina, Bonny Koane, David C. Bardos, Samuel Jeppy, and Vojtech Novotny. 2019. ‘Species Richness of Birds along a Complete Rain Forest Elevational Gradient in the Tropics: Habitat Complexity and Food Resources Matter’. Journal of Biogeography 46 (2): 279–90. 10.1111/jbi.13482.

Sayol, Ferran, Bouwe R. Reijenga, Joseph A. Tobias, and Alex L. Pigot. 2025. ‘Ecophysical Constraints on Avian Adaptation and Diversification’. Current Biology 35 (6): 1326–36.

Schumm, Matthew, Alexander E. White, K. Supriya, and Trevor D. Price. 2020. ‘Ecological Limits as the Driver of Bird Species Richness Patterns along the East Himalayan Elevational Gradient’. The American Naturalist 195 (5): 802–17. 10.1086/707665.

Seeholzer, Glenn F., Santiago Claramunt, and Robb T. Brumfield. 2017. ‘Niche Evolution and Diversification in a Neotropical Radiation of Birds (Aves: Furnariidae)’. Evolution 71 (3): 702–15. 10.1111/evo.13177.

Sheldon, Kimberly S. 2019. ‘Climate Change in the Tropics: Ecological and Evolutionary Responses at Low Latitudes’. Annual Review of Ecology, Evolution, and Systematics 50 (1): 303–33. 10.1146/annurev-ecolsys-110218-025005.

Sherry, Thomas W. 1990. ‘When Are Birds Dietarily Specialized? Distinguishing Ecological from Evolutionary Approaches’. Studies in Avian Biology 13: 337–52.

Sherry, Thomas W, Cody M Kent, Natalie V Sánchez, and Çağan H Şekercioğlu. 2020. ‘Insectivorous Birds in the Neotropics: Ecological Radiations, Specialization, and Coexistence in Species-Rich Communities’. The Auk 137 (4): 1–27. 10.1093/auk/ukaa049.

Sherry, Thomas W., and Lucinda A. McDade. 1982. ‘Prey Selection and Handling in Two Neotropical Hover-Gleaning Birds’. Ecology 63 (4): 1016–28. 10.2307/1937241.

Sillett, T. Scott, Anne James, and Kristine B. Sillett. 1997. ‘Bromeliad Foraging Specialization and Diet Selection of Pseudocolaptes Lawrencii (Furnariidae)’. Ornithological Monographs, 733–42.

Smith, Brian Tilston, John E. McCormack, Andrés M. Cuervo, et al. 2014. ‘The Drivers of Tropical Speciation’. Nature 515 (7527): 7527. 10.1038/nature13687.

Srinivasan, Umesh, Kartik Shanker, and Trevor D. Price. 2024. ‘Ant Impacts on Global Patterns of Bird Elevational Diversity’. Ecology Letters 27 (8): e14497. 10.1111/ele.14497.

Supriya, K., Corrie S. Moreau, Katerina Sam, and Trevor D. Price. 2019. ‘Analysis of Tropical and Temperate Elevational Gradients in Arthropod Abundance’. Frontiers of Biogeography 11 (2). 10.21425/F5FBG43104.

Terborgh, John W., and John Faaborg. 1980. ‘Saturation of Bird Communities in the West Indies’. The American Naturalist 116 (2): 178–95.

Thormann, Birthe, Dirk Ahrens, Carlos Iván Espinosa, et al. 2018. ‘Small-Scale Topography Modulates Elevational α-, β- and γ-Diversity of Andean Leaf Beetles’. Oecologia 187 (1): 181–89. 10.1007/s00442-018-4108-4.

Tobias, Joseph A., Catherine Sheard, Alex L. Pigot, et al. 2022. ‘AVONET: Morphological, Ecological and Geographical Data for All Birds’. Ecology Letters 25 (3): 581–97. 10.1111/ele.13898.

Winger, Benjamin M., and John M. Bates. 2015. ‘The Tempo of Trait Divergence in Geographic Isolation: Avian Speciation across the Marañon Valley of Peru’. Evolution 69 (3): 772–87. 10.1111/evo.12607.

Zaccarelli, Nicola, Daniel I. Bolnick, and Giorgio Mancinelli. 2013. ‘RInSp: An r Package for the Analysis of Individual Specialization in Resource Use’. Methods in Ecology and Evolution 4 (11): 1018–23. 10.1111/2041-210X.12079.

