## Supplemental Material for "Foraging behavior, not prey identity, facilitates niche packing in a tropical montane avifauna"

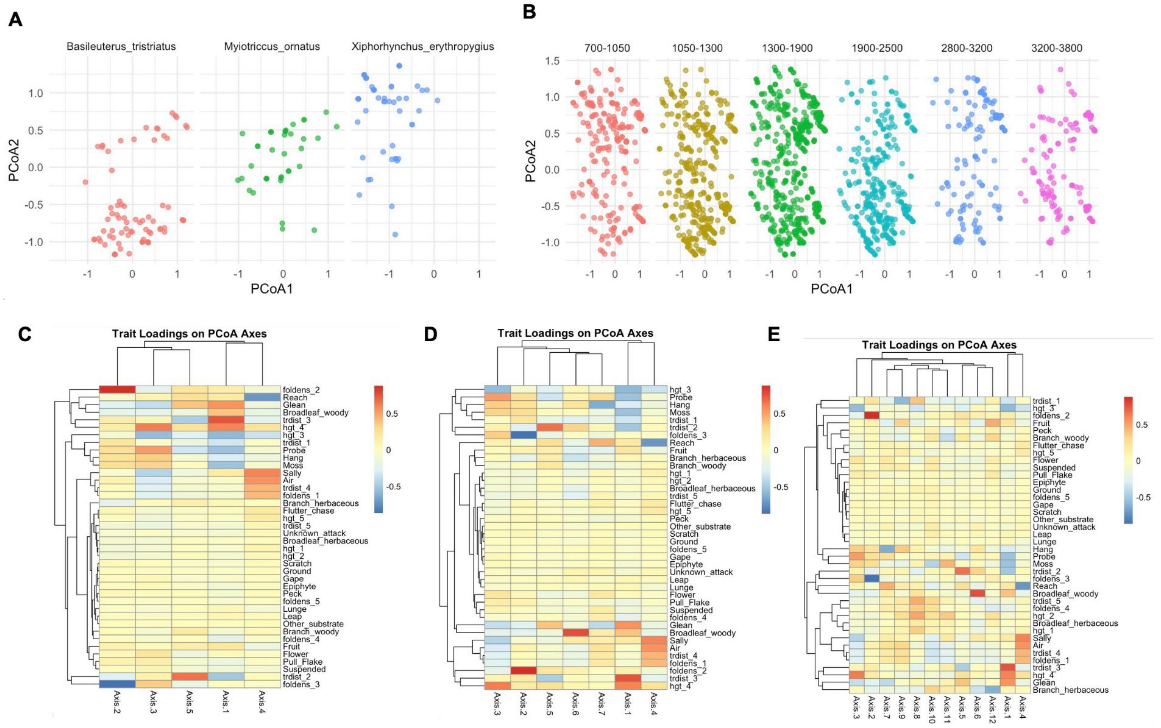

**Figure S1: Principal Coordinate Analysis of Foraging Behavior**
Panels show spread of foraging behavior in ordinated space for (A) three ecologically distinct species and (B) across six elevational bins, instead of the four that we use throughout the paper, with each point corresponding to a single observation. C-E show heatmaps and corresponding dendrograms of the positive and negative strength with which each behavior loads onto each axis, shown separately when considering five axes (54.17% total variance), seven axes (65.67% total variance) and twelve axes (80.85% total variance).

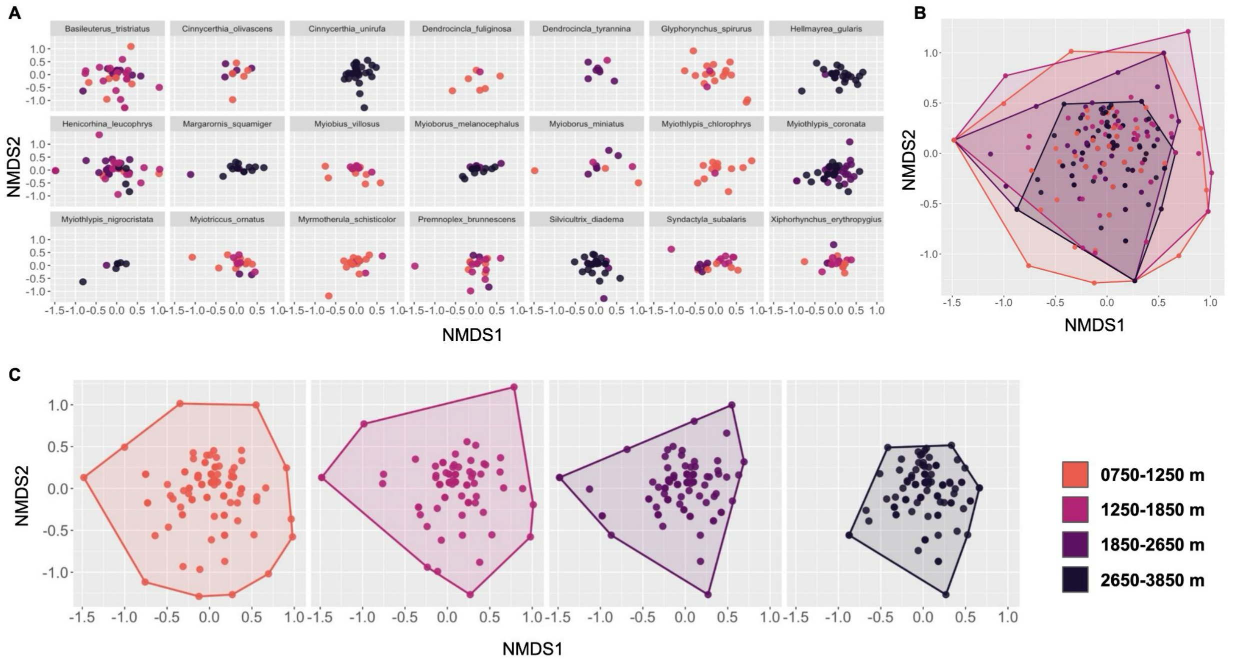

**Figure S2: NMDS of prey orders across elevational bins**
Non-metric multidimensional scaling shows that composition of invertebrate prey community orders are nested across elevation, with high-elevation communities representing a subset of low-elevation communities. Each point corresponds to an individual fecal sample, with convex hulls drawn around their extent for each elevational bin to visualize nestedness (NODF = 50.4, Matrix fill = 0.6, Stress = 0.18). Plotted separately for A) species, B) all elevational bins overlaid, and C) each elevational bin.

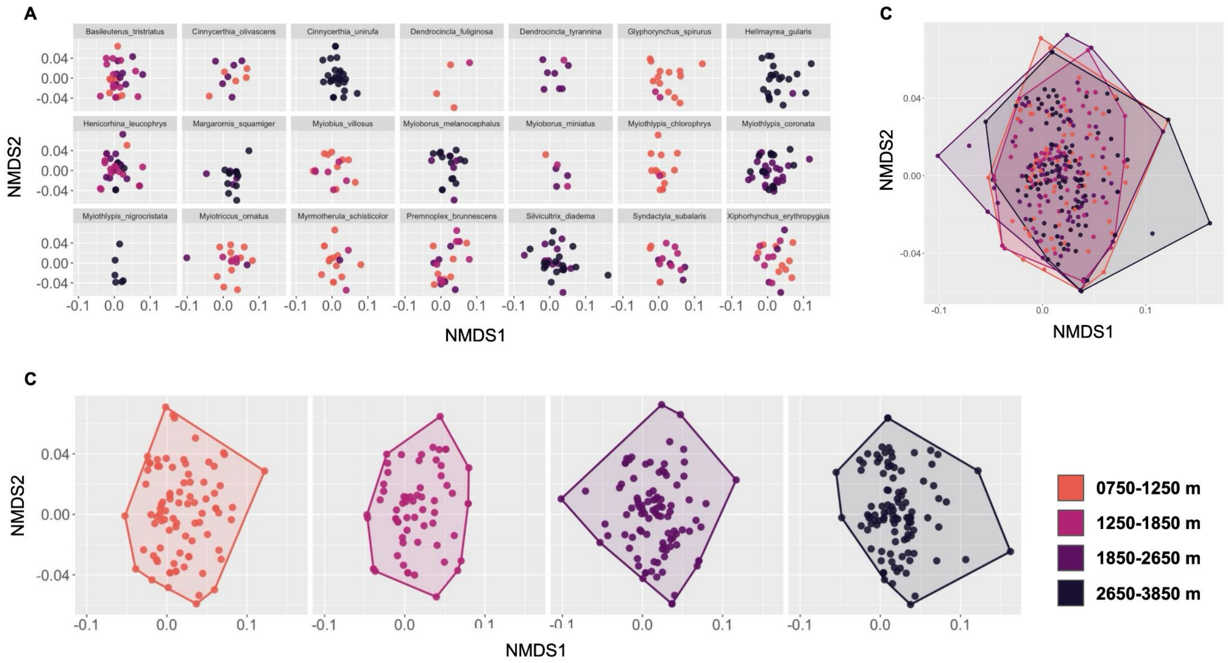

**Figure S2: NMDS of prey families across elevational bins**
Non-metric multidimensional scaling shows that composition of invertebrate prey community families are nested across elevation, with high-elevation communities representing a subset of low-elevation communities. Each point corresponds to an individual fecal sample, with convex hulls drawn around their extent for each elevational bin to visualize nestedness (NODF = 36.4, Matrix fill = 0.4, Stress = 0.14). Plotted separately for A) species, B) all elevational bins overlaid, and C) each elevational bin.

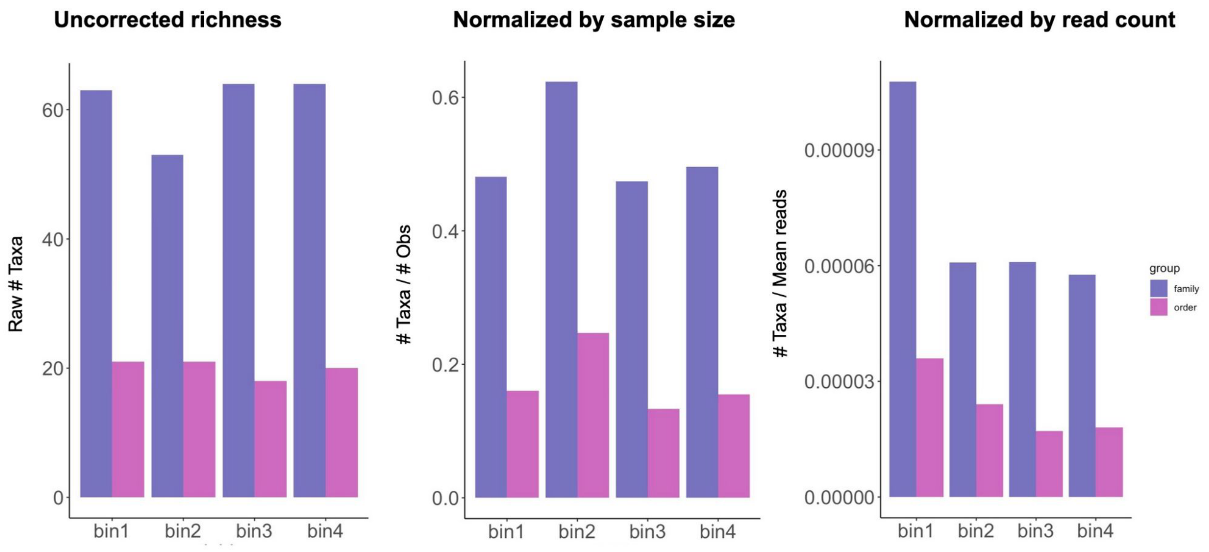

**Figure S4: Bar plots of prey counts**
Counts of invertebrate prey orders and families in each of four elevation bins, A) uncorrected, B) normalized by the number of samples in each elevational bin, and C) normalized by the mean number of reads in each elevational bin. Bin 1 = 750 – 1250 m, Bin 2 = 1250 – 1850 m, Bin 3 = 1850 – 2650 m, Bin 4 = 2650 – 3850 m.

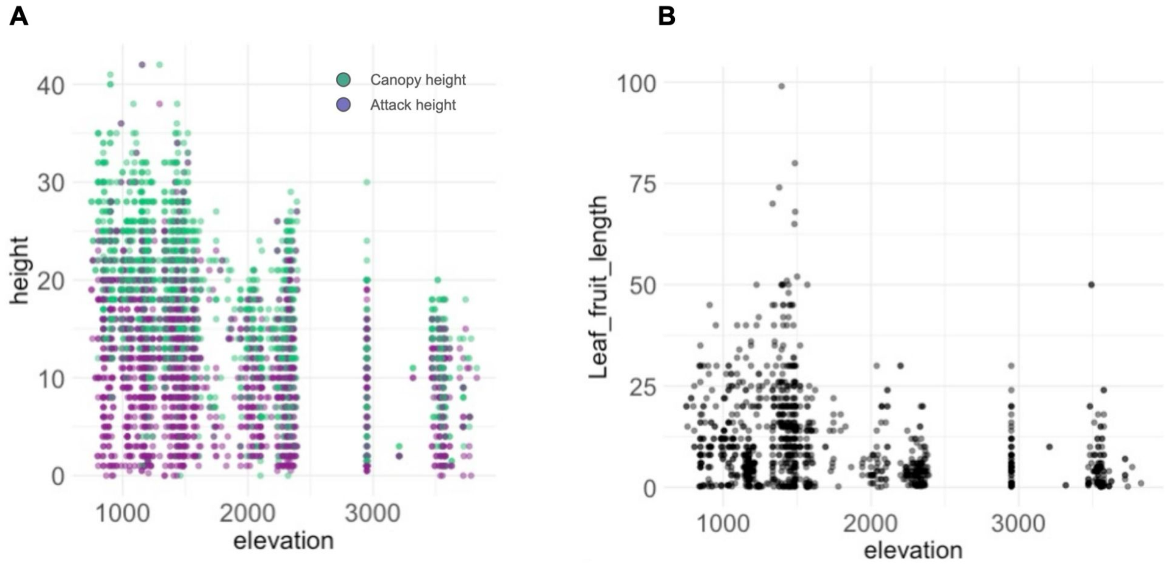

**Figure S5: Vegetation structure across elevation**
Change in forest structure across elevation based on visual estimates of A) canopy and foraging height and B) the length of the leaf, fruit, or flower where a foraging maneuver occurred.

**Table S1: Summary of bird species with metabarcoded fecal samples**

| Scientific name | N obs | Mean reads | Total reads | N orders | N families | Elev midpoint | Elev range |
|---|---|---|---|---|---|---|---|
| Basileuterus_tristriatus | 42 | 553958.81 | 23266270 | 12 | 41 | 1600 | 1500 |
| Cinnycerthia_olivascens | 12 | 920569.917 | 11046839 | 12 | 27 | 1650 | 1400 |
| Cinnycerthia_unirufa | 30 | 1585081.5 | 47552445 | 18 | 50 | 3200 | 1300 |
| Dendrocincla_fuliginosa | 8 | 685222.125 | 5481777 | 8 | 14 | 900 | 1500 |
| Dendrocincla_tyrannina | 8 | 5067500 | 40540000 | 11 | 24 | 2250 | 1700 |
| Glyphorynchus_spirurus | 19 | 370679 | 7042901 | 18 | 35 | 1000 | 1700 |
| Hellmayrea_gularis | 24 | 685338.75 | 16448130 | 13 | 37 | 3200 | 1300 |
| Henicorhina_leucophrys | 48 | 737165 | 35383920 | 19 | 50 | 1950 | 2400 |
| Margarornis_squamiger | 19 | 2066674.42 | 39266814 | 13 | 39 | 2950 | 2000 |
| Myiobius_villosus | 18 | 901825.278 | 16232855 | 11 | 37 | 1150 | 1200 |
| Myioborus_melanocephalus | 21 | 687488.619 | 14437261 | 10 | 40 | 3050 | 1600 |
| Myioborus_miniatus | 11 | 286534.273 | 3151877 | 11 | 20 | 1500 | 2100 |
| Myiothlypis_chlorophrys | 18 | 313987.444 | 5651774 | 12 | 26 | 950 | 800 |
| Myiothlypis_coronata | 51 | 546842.373 | 27888961 | 15 | 48 | 2450 | 1600 |
| Myiothlypis_nigrocristata | 7 | 215213.714 | 1506496 | 8 | 17 | 2950 | 1400 |
| Myiotriccus_ornatus | 20 | 1193998 | 23879960 | 15 | 43 | 1200 | 1700 |
| Myrmotherula_schisticolor | 22 | 818310.818 | 18002838 | 13 | 32 | 1000 | 1700 |
| Premnoplex_brunnescens | 24 | 1218046.25 | 29233110 | 13 | 30 | 1450 | 1200 |
| Pseudotriccus_pelzelni | 1 | 430596 | 430596 | 8 | 5 | 1400 | 1500 |
| Silvicultrix_diadema | 31 | 1265975.55 | 39245242 | 18 | 48 | 2650 | 1000 |
| Syndactyla_subalaris | 21 | 415761.667 | 8730995 | 17 | 33 | 1300 | 2100 |
| Xiphorhynchus_erythropygius | 25 | 851876.28 | 21296907 | 14 | 34 | 1000 | 1700 |

**Table S2: Counts of detected invertebrate prey orders and families across fecal samples**

| order | # of samples observed |
|---|---|
| Lepidoptera | 423 |
| Diptera | 421 |
| Hemiptera | 367 |
| Araneae | 358 |
| Coleoptera | 332 |
| Isopoda | 313 |
| Hymenoptera | 247 |
| Orthoptera | 74 |
| Blattodea | 66 |
| Haplotaxida | 17 |
| Entomobryomorpha | 14 |
| Ixodida | 11 |
| Opiliones | 11 |
| Trombidiformes | 10 |
| Neuroptera | 8 |
| Psocodea | 8 |
| Stylommatophora | 6 |
| Cyclopoida | 4 |
| Ephemeroptera | 2 |
| Phasmatodea | 2 |
| Pseudoscorpiones | 2 |
| Enchytraeida | 1 |
| Lithobiomorpha | 1 |
| Plecoptera | 1 |

| | |
|---|---|
| Poduromorpha | 1 |
| Scolopendromorpha | 1 |
| Symphypleona | 1 |
| Thysanoptera | 1 |
| Trichoptera | 1 |
| **Family** | **# Of samples observed** |
| Geometridae | 214 |
| Anyphaenidae | 173 |
| Curculionidae | 79 |
| Ichneumonidae | 79 |
| Braconidae | 76 |
| Chrysomelidae | 72 |
| Salticidae | 68 |
| Miridae | 66 |
| Sciaridae | 63 |
| Erebidae | 54 |
| Ectobiidae | 50 |
| Nymphalidae | 50 |
| Formicidae | 47 |
| Cicadidae | 43 |
| Pompilidae | 36 |
| Tachinidae | 35 |
| Limoniidae | 33 |
| Tipulidae | 25 |
| Oedemeridae | 24 |
| Syrphidae | 21 |

| | |
|---|---|
| Elateridae | 20 |
| Theridiidae | 19 |
| Ptilodactylidae | 18 |
| Muscidae | 16 |
| Mycetophilidae | 15 |
| Cecidomyiidae | 13 |
| Staphylinidae | 13 |
| Cicadellidae | 12 |
| Ceratopogonidae | 11 |
| Scarabaeidae | 11 |
| Tabanidae | 11 |
| Chironomidae | 10 |
| Linyphiidae | 10 |
| Stratiomyidae | 9 |
| Carabidae | 8 |
| Psychodidae | 7 |
| Pyralidae | 7 |
| Apidae | 6 |
| Culicidae | 6 |
| Rhyparochromidae | 6 |
| Tetragnathidae | 6 |
| Crambidae | 5 |
| Drosophilidae | 5 |
| Anthomyzidae | 4 |
| Asilidae | 4 |
| Coccinellidae | 4 |
| Diapriidae | 4 |

| | |
|---|---|
| Hesperiidae | 4 |
| Pentatomidae | 4 |
| Aphididae | 3 |
| Bibionidae | 3 |
| Dryinidae | 3 |
| Fulgoridae | 3 |
| Ixodidae | 3 |
| Keroplatidae | 3 |
| Tenebrionidae | 3 |
| Tortricidae | 3 |
| Vespidae | 3 |
| Bethylidae | 2 |
| Flatidae | 2 |
| Halictidae | 2 |
| Nitidulidae | 2 |
| Nolidae | 2 |
| Phasmatidae | 2 |
| Riodinidae | 2 |
| Sarcophagidae | 2 |
| Thomisidae | 2 |
| Tydeidae | 2 |
| Achilidae | 1 |
| Aleyrodidae | 1 |
| Anthribidae | 1 |
| Blattidae | 1 |
| Calamoceratidae | 1 |
| Cerambycidae | 1 |

| | |
|---|---|
| Chloropidae | 1 |
| Chrysopidae | 1 |
| Colletidae | 1 |
| Cosmetidae | 1 |
| Cyrtaucheniidae | 1 |
| Delphacidae | 1 |
| Encyrtidae | 1 |
| Erotylidae | 1 |
| Euconulidae | 1 |
| Evaniidae | 1 |
| Figitidae | 1 |
| Gryllidae | 1 |
| Hahniidae | 1 |
| Hemerobiidae | 1 |
| Hybotidae | 1 |
| Lycidae | 1 |
| Megalopygidae | 1 |
| Noctuidae | 1 |
| Notodontidae | 1 |
| Pemphredonidae | 1 |
| Phalacridae | 1 |
| Phoridae | 1 |
| Platygastridae | 1 |
| Scatopsidae | 1 |
| Scirtidae | 1 |
| Scolopocryptopidae | 1 |
| Tarsonemidae | 1 |

| | |
|---|---|
| Tineidae | 1 |
| Tiphiidae | 1 |
| Trigonidiidae | 1 |
| Xylomyidae | 1 |
